# Intra-Strain Genetic Heterogeneity in *Toxoplasma gondii* ME49: Oxford Nanopore Long-Read Sequencing Reveals Copy Number Variation in the ROP8-ROP2A Locus

**DOI:** 10.1101/2025.05.10.653050

**Authors:** Yomna Gohar, Marie Neumann, Lisanna Hülse, Daniel Wind, Julia Mock, Karin Buchholz, Marcel Helle, Ursula R Sorg, Daniel Degrandi, Klaus Pfeffer, Alexander Dilthey

## Abstract

**Background:** *Toxoplasma gondii* is an important pathogen and model organism for studying mechanisms of immune evasion and defense. Within the same strain, model organisms are typically assumed to be isogenic; for *T. gondii*, within-strain genetic divergence has been detected based on phenotypic changes and older molecular techniques but not characterized at the genomic level. We therefore used Oxford Nanopore long-read sequencing to characterize three independently maintained *T. gondii* ME49 isolates: 2015T and 2020T (obtained from ATCC and propagated in cell culture), and 2000B (propagated in mice).

**Results:** We *de novo* assembled a new *T. gondii* ME49 reference genome and, using state of-the-art variant calling combined with pangenomic genotyping, detected variants between the sequenced isolates. Our new reference genome exceeded existing reference genomes in continuity (NG50 = 6.68 Mb versus 1.2 Mb in RefSeq) and structural accuracy, resolving all chromosomes except for a single break in the ribosomal DNA region. For isolates 2000B and 2020T, we identified 106 and 128 variants, respectively, across a final call set of 79 SNVs, 93 INDELs, and five structural variants; 18 small non-synonymous variants included genes associated with *T. gondii* life cycle (AP2X-8) and virulence *in vivo* (6-phosphogluconate dehydrogenase). A 13 kb expansion in the ROP8-ROP2A virulence locus increased the copy number of ROP2A-ROP8 genes in isolates 2000B and 2020T from three to six.

**Conclusions:** We provide an improved *T. gondii* ME49 reference genome and demonstrate the potentially confounding effect of intra-strain genetic heterogeneity, highlighting the need for continuous genomic monitoring for long-term genetic identity.

## Background

Model organisms are essential in biological research, providing controlled systems to study fundamental biological processes. A key assumption is that representatives of a given strain are largely isogenic. However, prolonged laboratory maintenance, for example after acquisition from a commercial vendor (e.g., ATCC) or a partner lab, can lead to genetic divergence, which — unless it has obvious phenotypic effects—often goes undetected [1–4]. Studies have demonstrated that genetic heterogeneity accumulates during laboratory maintenance and impacts research outcomes and phenotypic variation in key model organisms, including laboratory mice (*Mus musculus*) [5–8], *Drosophila melanogaster* [9], *Aedes aegypti* (the yellow fever mosquito) [10], bacteria [11–13], *C. elegan*s [14] and different human cell lines inculding human pluripotent stem cells (hPSCs) [15–18], which serve as in vitro model systems for studying human biology and disease. Genetic heterogeneity can encompass several types of variation, including single nucleotide polymorphisms (SNPs), small insertions and deletions (INDELs), and structural variants (SVs), the latter of which is often difficult to detect [8,19,20]. Mechanisms for the generation of genetic heterogeneity include replication errors, DNA repair mechanisms, and selection pressures [21–23]. In vivo passaging, in particular, exposes strains to host immune responses, accelerating genetic drift and selection [24,25].

Among microbial pathogens, *Toxoplasma gondii* serves as a crucial model organism for studying host-pathogen interactions, immune evasion, and virulence mechanisms [26,27]. Most *T. gondii* strains found in North America and Europe belong to three clonal lineages—types I, II, and III [28]. Type II strains are responsible for most of the human infections studied in North America and Europe, and are likewise prevalent in livestock from these regions [28]. Previous studies on various *T. gondii* strains have shown that continuous passaging in mice or cell culture can lead to genetic and phenotypic changes. For example, research on Type I RH-derived clonal lineages, using restriction fragment length polymorphism (RFLP) analysis, has demonstrated that repeated passaging could induce genomic heterogeneity [29]. Some RH-derived lineages have also showed phenotypic divergence, such as variation in plaque size, growth rate, differentiation, and ability to survive outside host cells [30]. Additionally, prolonged laboratory maintenance of *T. gondii* through serial passaging in mice or cell culture has been linked to the loss of oocyst production in cats, as observed in multiple strains, including M-7741, GT-1, and RH, likely due to accumulated genetic changes over time [31–33].

While these studies have demonstrated that laboratory propagation of *T. gondii* can induce genetic heterogeneity associated with phenotypic changes, they were conducted before the emergence of high-throughput sequencing and did not include a characterization of genetic heterogeneity at the sequence level. What is more, comparative genomic studies in the *T. gondii* field in general have traditionally relied on short-read sequencing technologies [34,35]. Short-read sequencing technologies have inherent limitations in detecting complex structural variations, as short reads cannot span large genomic rearrangements, repetitive regions, or highly polymorphic loci [36]. As a result, the importance of complex genetic variation to overall *T. gondii* genetic diversity remains incompletely characterized.

In contrast, long-read sequencing technologies, such as Oxford Nanopore sequencing, can overcome these challenges by producing long continuous reads, enabling more comprehensive detection of genomic variants. Recent advances, particularly in the Oxford Nanopore R10.4 platform, have further improved sequencing accuracy, allowing the generation of near-finished microbial genomes without the need for short-read or reference-based polishing [37].

In this study (Figure 1), we leveraged Oxford Nanopore long-read sequencing to assess genomic divergence among three independently maintained *T. gondii* ME49 isolates—a widely used type II reference strain with moderate virulence [38,39]. Employing de novo assembly and a multimethod variant calling approach, followed by graph genome-based genotyping (Supplementary Figure S1), our analysis revealed genetic heterogeneity among these isolates, characterized by both small variants and large structural variations that would have been inaccessible using traditional sequencing methods. Our findings contribute to a broader understanding of *T. gondii* genomic variation, with implications for strain characterization and experimental reproducibility.

**Figure 1:**
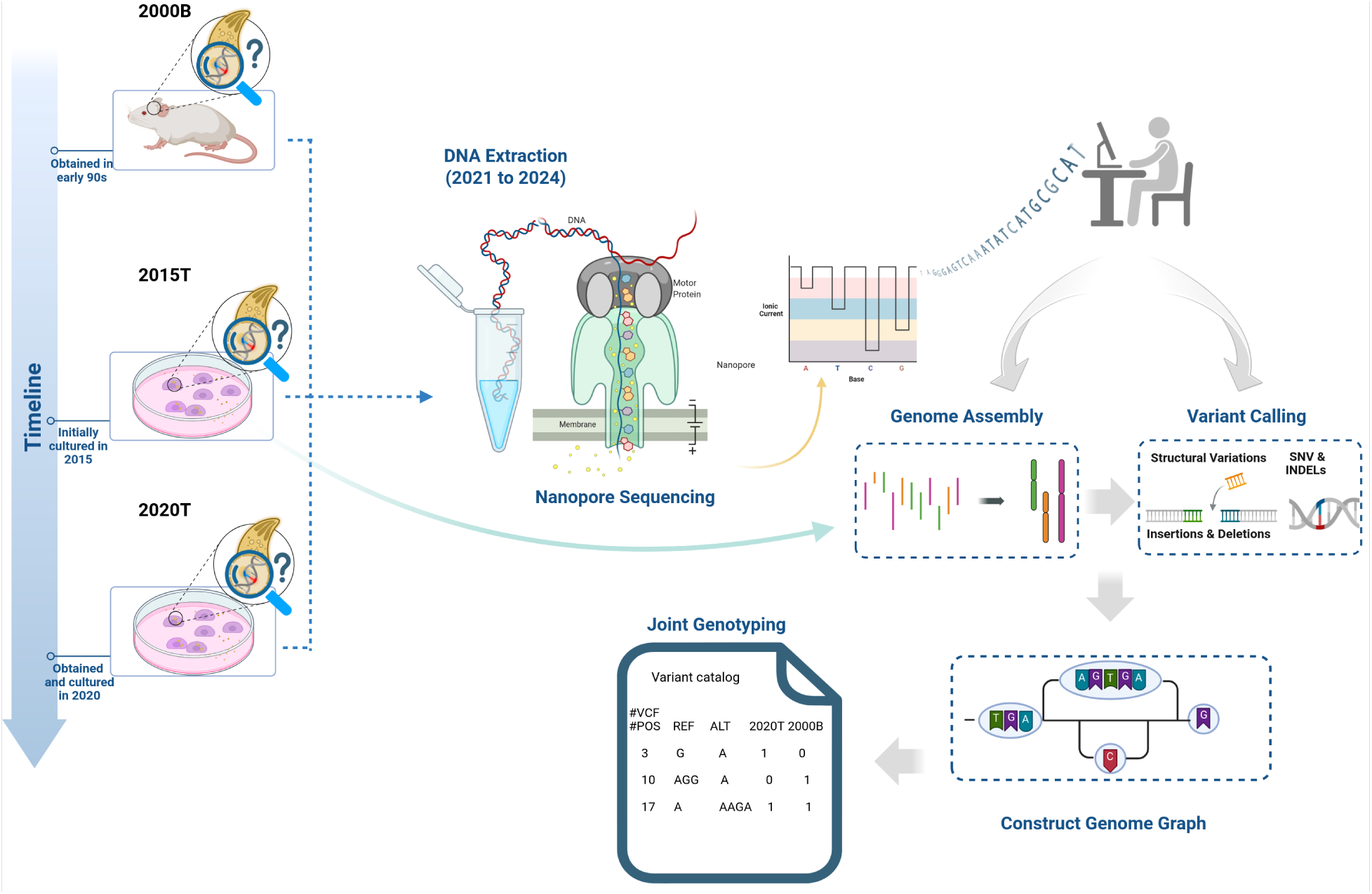
Overview of the sequencing and variant analysis pipeline for *T. gondii* ME49 isolates. Created in BioRender. https://BioRender.com/undefined

## Results

### Generation of Nanopore sequencing data for three *T. gondii* ME49 isolates

We selected three ME49 *T. gondii* isolates for characterization: 2015T, obtained from ATCC (catalog number 50611) thawed in 2015, expanded for two passages, refrozen and then continuously propagated in cell culture from 2022 onward; 2020T, obtained in 2020 from ATCC (catalog number 50611) and propagated in cell culture; and 2000B, originally obtained in the early 1990s and propagated in CD1 mice (Figure 2A; see Supplementary Note for additional details on the propagation history of these isolates). We generated single-molecule long-read Oxford Nanopore R10 sequencing data for the three isolates (Figure 2B), yielding 203X genomic coverage at a read N50 of 37kb for isolate 2015T; 20X genomic coverage at a read N50 of 5kb for isolate 2020T; and 23X genomic coverage at a read N50 of 12.8kb for isolate 2000B. For isolate 2015T, all sequencing data was generated with the R10.4.1 sequencing chemistry and basecalling was carried out with Dorado [40] in “super high accuracy” mode, followed by HERRO (haplotype-aware error correction of ultra-long nanopore reads) [41] for error correction; for the other isolates, multiple sequencing chemistries in the R10 family were employed and basecalling was carried out with Guppy [42] in “super high accuracy” mode (see Supplementary Table 1 for details on the generated sequencing data, proportion of host DNA, sequencing chemistries and basecalling). Preliminary analyses of the generated data indicated that reads containing mitochondrial DNA fragments led to challenges during further data analyses; such reads were therefore removed (Supplementary Notes).

**Figure 2.**
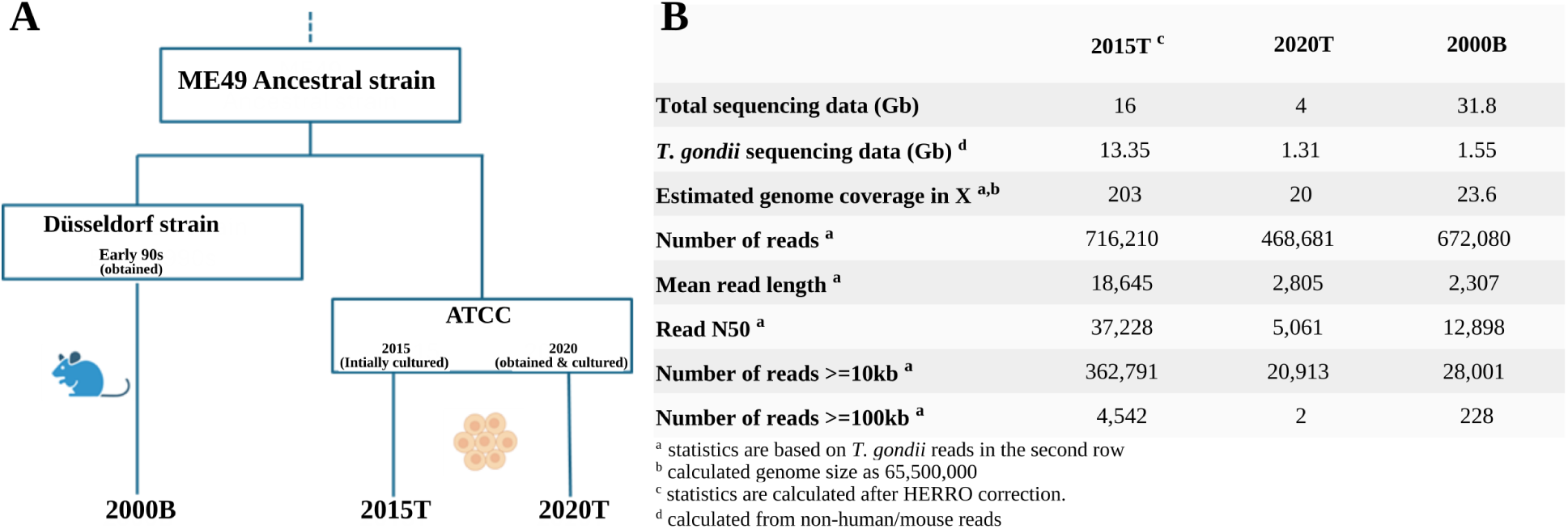
Origin and sequencing summary of *Toxoplasma gondii* ME49 isolates used in this study. (A) Schematic overview of the propagation and acquisition history of the ME49 isolates 2000B, 2015T, and 2020T. (B) Summary statistics for Oxford Nanopore long-read sequencing data generated for each isolate.

### High-quality genome assembly for isolate 2015T

We assembled the high-quality HERRO-corrected sequencing data for isolate 2015T using Flye [43] and obtained a genome assembly consisting of 22 contigs and a total size of 64.5 Mb at an NG50 of 6.68Mb (Figure 3A). We investigated the chromosomal completeness of our assembly and found that 12 of 13 *T. gondii* ME49 chromosomes were almost completely covered by single assembly contigs; the remaining chromosome IX was split into two contigs (Figure 3B, 3C). The genome of the *T. gondii* ME49 apicoplast, which is approximately 35kb in size and particularly challenging to assemble due to the presence of inverted repeats [44], was represented by two contigs of approximately 23 and 5 kb in size. We annotated our assembly with Companion [45] and found that it contained 9,137 genes and 166 pseudogenes.

**Figure 3.**
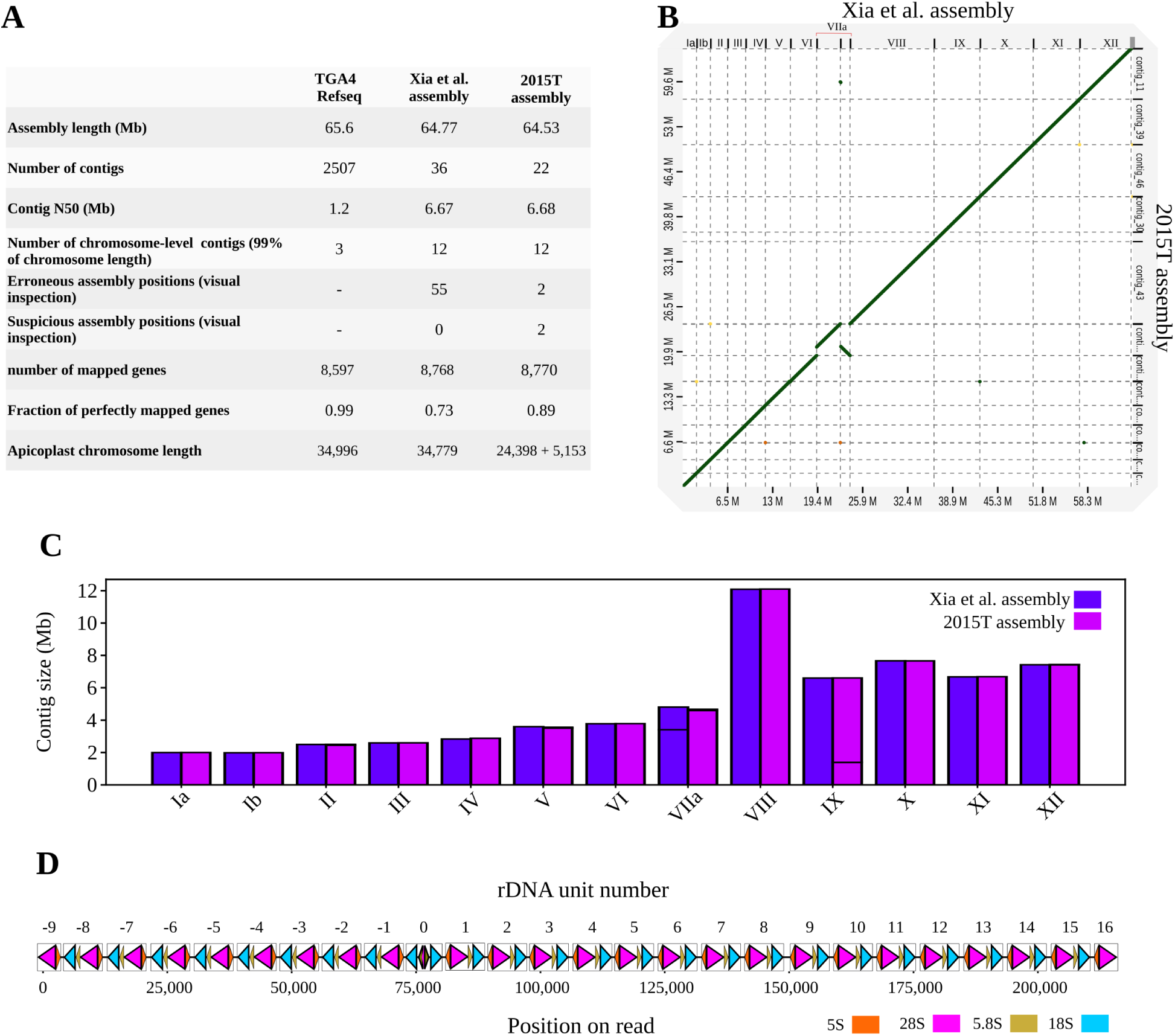
Comparison of the 2015T assembly to existing *T. gondii* ME49 references and detailed view of the rDNA locus. (A) Comparison of the 2015T genome assembly to previously published *T. gondii* ME49 reference genomes. (B) Dot plot showing structural alignment between the 2015T and Xia et al. Assemblies. (C) Contig size comparison, with horizontal lines indicating breaks between contigs. (D) Partial sequence structure of the ribosomal DNA (rDNA) locus in isolate 2015T, visualized using the longest read aligned to this region. Triangles mark the positions of 18S, 28S, 5.8S, and 5S rRNA genes along the read. The orientation of each triangle indicates the direction of rRNA gene alignment: left-facing triangles represent reverse-strand mappings, and right-facing triangles represent forward-strand mappings.

### Comparison of 2015T to other existing reference assemblies

To evaluate the quality of our assembly, we compared it to two existing *T. gondii* ME49 reference genomes (Figure 3A), the TGA4 *T. gondii* reference genome from RefSeq and a more recent single-molecule long-read sequencing-based assembly [46], referred to as the “Xia et al. assembly” for the remainder of this paper. Compared to TGA4 (NCBI RefSeq assembly: GCF_000006565.2), our assembly was much more contiguous (NG50 of 6.68Mb for our assembly compared to 1.2Mb for TGA4); in addition, TGA4 was reported to contain unresolved duplications (e.g., in the ROP4/ROP7 locus) and an erroneous inversion on chromosome 4 [46]. The Xia et al. assembly exhibited comparable contiguity to our assembly (36 contigs at a NG50 of 6.67Mb); we thus carried out an in-depth comparison. First, we carried out a dot plot analysis which revealed a high degree of collinearity between Xia et al. and our assembly, with few deviations observed for Chromosomes VIIa and IX (Figure 3B). Chromosome VIIa was split in two fragments of approximately 3.4Mb and 1.3Mb in size in Xia et al., whereas our assembly contained a near-complete chromosome VIIa contig of 4.5Mb in size. Chromosome IX was split into two contigs in our assembly, whereas in Xia et al. it was represented by a single contig.

Otherwise, both assemblies exhibited almost identical contig sizes (Figure 3C). Second, we investigated the size differences between the two assemblies. Using nucmer [47], we found that each assembly contained a stretch of sequence nearly identical in length, that could not be aligned to the other assembly (76kb for Xia et al. compared to our assembly and 86kb for our assembly compared to Xia et al.), suggesting that both assemblies covered comparable fractions of the *T. gondii* ME49 genome despite the slightly increased size of the Xia et al. assembly.

Third, the proportion of *T. gondii* gene sequences from ToxoDB [48] with perfect alignments was higher for our assembly than for Xia et al. (89% compared to 73%, see Figure 3A and Figure S2 for examples of genes exhibiting more accurate mapping in our assembly), and annotation of the Xia et al. assembly using the same Companion approach we applied to our 2015T assembly revealed a higher fraction of pseudogenes (8891 genes and 557 pseudogenes). These results indicated a higher consensus quality of our assembly compared to the Xia et al. assembly, likely associated with the higher error rate of older Nanopore sequencing data used by Xia et al. (MinION R9.4.1).

Fourth, manual inspection of read alignments (based on the reads used by Xia et al. for assembly construction) in Integrative Genomic Viewer (IGV) [49] identified 55 regions in the Xia et al. assembly that were likely structurally incorrect (Supplementary Table 2, Supplementary Notes). We projected the coordinates of these onto our assembly and, based on visual assessment in IGV, found that all but 4 of the evaluated positions were correctly resolved in our assembly. One of the 4 unresolved positions coincided with the Chromosome IX breakpoint (see above); visual inspection in Xia et al. was indicative of a misassembled repetitive region with a significant read depth increase and numerous reads with a mapping quality of zero at that position (Figure S3). Analysis of reads from the region using BLAST [50] indicated the presence of ribosomal DNA (rDNA) sequences. We extracted the longest read (207,813 bp) from our 2015T dataset that mapped to the rDNA locus in the Xia et al. assembly and found that it contained at least 23 blocks of repetitive rDNA units (Figure 3D). Notably, the read indicated a shift in relative rDNA repeat unit orientation in the middle of the rDNA region captured by the read, which may contribute to the assembly challenges at this locus. In contrast, the corresponding position in the Xia et al. assembly showed only 16 rDNA blocks (Figure S3). Together, the discrepancy in rDNA copy number and the visual evidence from IGV suggested that, despite the representation of Chromosome IX as a single contig in the Xia et al. assembly, the assembly of this region in Xia et al. was not accurate.

We thus concluded that our assembly was - with the possible exception of the apicoplast genome, which was represented as a single contig of approximately 35kb in size in the Xia et al. assembly (Figure S4) - of higher quality than the Xia et al. assembly, likely reflecting the markedly higher coverage of the dataset used to construct our assembly (203X vs 30X) and the improved quality of R10.4.1 sequencing data. A final visual inspection of the complete assembly confirmed the structural accuracy of our assembly, with the possible exception of telomeric regions, for which we did not carry out an in-depth assessment.

### Variant discovery in isolates 2020T and 2000B

To detect candidate polymorphisms between the three isolates sequenced by in this study, we carried out variant calling for isolates 2020T and 2000B against the 2015T assembly, supplemented with the apicoplast sequence from Xia et al. to prevent read misalignment. For discovery of candidate small variants, i.e. of single nucleotide polymorphisms (SNPs) and small insertions/deletions (INDELs), we used medaka [51]. To minimize false-positive calls, we applied stringent filtering criteria (DP ≥ 10; GQ ≥ 10; removal of variants from structurally anomalous regions; see Materials and Methods), reducing the total set of small variant calls from 4,076 to 365 in sample 2020T and from 2,041 to 376 in 2000B (Supplementary Table 3). The post-filtering candidate variant set contained 67 SNPs and 298 INDELs for 2020T and 48 SNPs and 328 INDELs for 2000B (Supplementary Table 3). We observed a strong enrichment of candidate variant calls in repetitive and homopolymer regions (e.g., for both 2020T and 2000B about 97% of INDELs were in tandem repeat or homopolymeric regions; see Supplementary Table 4), suggesting the presence of false-positive calls despite the application of strict filtering criteria. For discovery of SVs, we employed Sniffles2 [52]. We performed manual inspection of SV candidates to remove false-positive calls in IGV (Materials and Methods), which reduced the total size of the Sniffles2-based SV call set from 116 to 2 for 2020T and from 36 to 3 in 2000B (Supplementary Table 5 and Supplementary Table 6). Post-filtering, we identified 367 small and large variants (8.7% of the total variants called) in 2020T and 379 variants (18% of the total variants called) in 2000B. Together, these variants comprised 693 unique variants across 669 unique loci, including SNPs, small indels, and structural variants.

### Discovery of a large structural variant in the ROP8-ROP2A locus by manual curation

During manual inspection of SV calls for isolate 2000B, we observed an enrichment of SV calls with low reported allele frequencies in the ROP8-ROP2A region on Chromosome X; while these calls were individually rejected during manual inspection, the pattern of the aligned reads, with the presence of >4,000bp of inserted sequence and exhibiting high frequencies of clipping and supplementary alignments, suggested the presence of a large structural variant that was not correctly resolved by Sniffles2 (Figure S5). Attempts at targeted re-assembly of the locus using standard assembly algorithms were not successful. However, we were able to manually assemble the locus by first determining its overall sequence structure based on overlaps between individual long reads (“read stitching”), followed by polishing (Supplementary Notes), revealing the presence of a 13 kb insertion. To confirm the existence of this 13 kb sequence in 2000B, we modified the 2015T assembly by incorporating the insertion using bcftools consensus. Alignment of 2000B reads to the modified reference showed a uniform coverage pattern without significant read clipping, showing that we had correctly resolved the structural variant (Figure S5). An initial inspection of the inserted sequence suggested that it contained additional copies of ROP8- and ROP2A-related genes (see below). Furthermore, visual inspection of the region in 2020T also indicated the likely presence of a structural variant relative to the 2015T assembly (Figure S6); however, due to the shorter read lengths of 2020T, we could not manually resolve this locus for 2020T. The 13 kb insertion (contig_46:7310767) from the stitched 2000B sequence was included in the SV variant set for subsequent joint genotyping.

To further characterize the sequence content of the inserted sequence in 2000B in comparison to the 2015T reference assembly, we carried out a fine-scale annotation effort, producing a set of consensus annotations based on manual curation of the output of three different tools (Augustus [53], Companion, BLAST; see Methods). We then compared the identified genes to the ME49 ROP2A and ROP8 sequences in ToxoDB and found that isolate 2000B contained three ROP2A-like genes and three ROP8-like genes, similar to the genome assembled by Xia et al., whereas 2015T had one ROP2A-like gene and two ROP8-like genes. Notably, the ME49 ROP2A sequence in ToxoDB is incomplete; gene classification in our assemblies was thus based on the shared sequence components of ROP2A and ROP8 reference genes. Accordingly, we refer to the ROP8-like sequences as ROP8-1, ROP8-2, and ROP8-3, and the ROP2A-like sequences as ROP2A-1, ROP2A-2A, and ROP2A-2B (Figure 4A).

**Figure 4.**
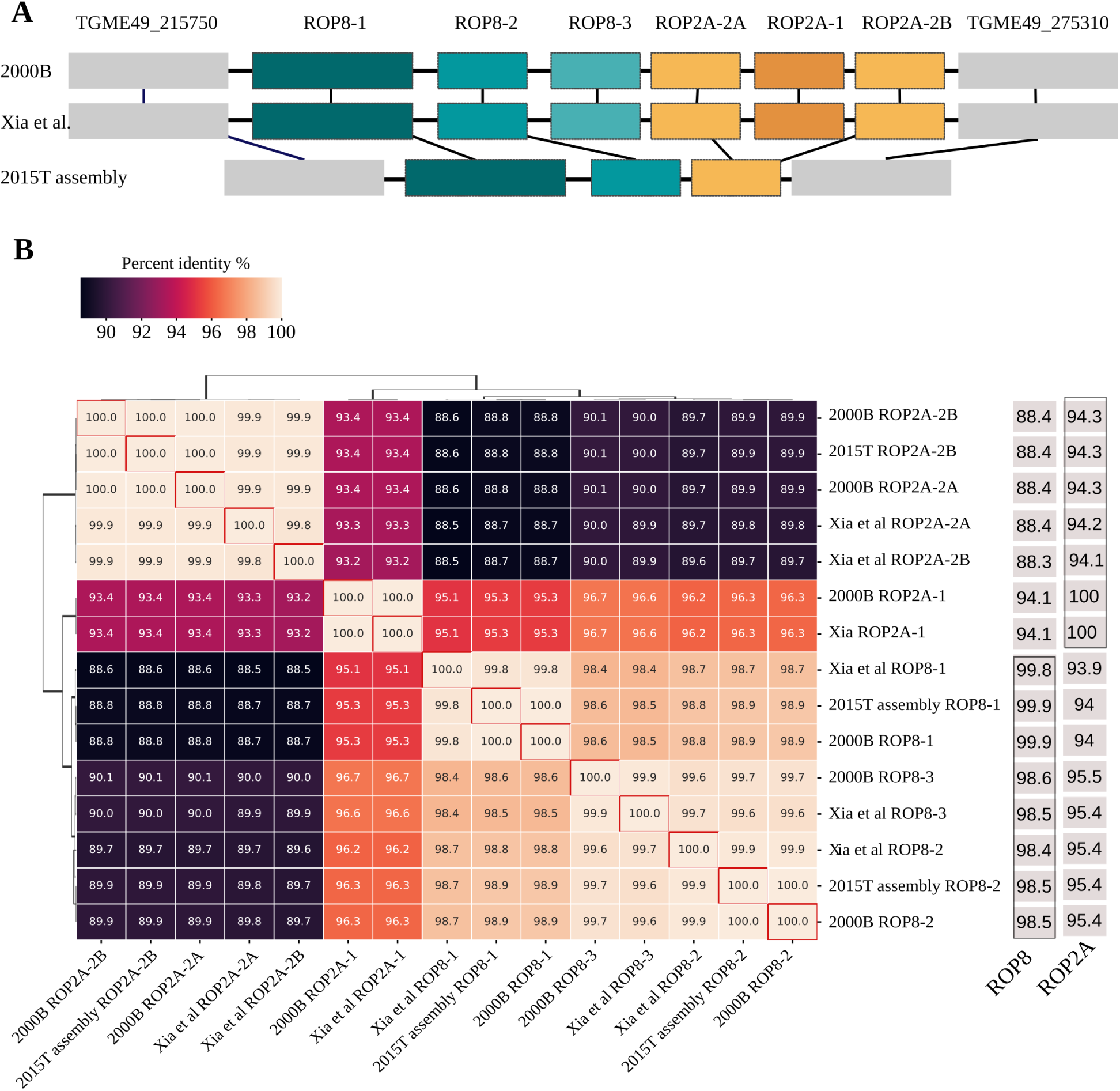
Structural and sequence variation in the ROP8–ROP2A locus. (A) Structural comparison of the ROP8–ROP2A locus across three assemblies: 2015T, 2000B, and the Xia et al. ME49 assembly. The copy number of ROP8 and ROP2A gene copies vary among isolates, with 2020T (not shown) carrying the same configuration as 2000B, as determined by graph-based genotyping. (B) Pairwise sequence identity heatmap of ROP8 and ROP2A gene copies across the 2015T, 2000B, and Xia et al. assemblies. Percent identity values are shown, with hierarchical clustering highlighting relationships among paralogs. Sequence identity to canonical ROP8 and ROP2A ME49 reference genes from ToxoDB is shown in the rightmost columns.

Furthermore, ROP8-1 and ROP8-2 were identical between isolates 2015T and 2000B, as was ROP2A-2B. ROP2A-2A was found only in 2000B, and its sequence was identical to ROP2A-2B (Figure 4B). Given that ROP8 and ROP2A are members of the ROP2 superfamily in *T. gondii*, our results suggest that the 2015T isolate may harbor three gene copies related to ROP8 and ROP2A, while both the 2000B isolate and the Xia et al. assembly contain six. These findings highlighted a substantial degree of copy number variation at a virulence-related locus of the *T. gondii* ME49 genome. To assess whether the additional gene copies were potentially functional, we mapped *T. gondii* ME49 RNA-seq data obtained from SRA (SRX6428507) to the resolved sequence of the SV from isolate 2000B and found evidence that ROP8-1, ROP8-2, ROP8-3, ROP2A-A, ROP2A-2B and ROP2A-1 were potentially expressed (Figure S8).

### Graph-based re-genotyping of 2015T, 2000B and 2020T

To establish a unified variant calling framework for the detection of high-quality isolate-distinguishing variants, we constructed a variation graph with vg [54], using the 2015T assembly as the reference and integrating the set of 670 small and large 2000B and 2020T candidate post-filtering variant loci. To produce graph-based variant calls, we re-aligned the generated Nanopore sequencing reads to the graph using GraphAligner [55], followed by re-genotyping with vg. For stringent quality control, we removed all positions for which the re-genotyping process did not yield a “reference” call for isolate 2015T, and all positions at which the frequency of the reference allele in isolate 2015T in the aligned reads was below 90%, leading to the removal of 487 loci. We found this step to be empirically necessary to reduce the rate of false-positive variant calls associated with homopolymer and tandem repeat regions, and with systematic differences of sequencing read quality between the sequenced isolates; of note, to reduce systematic differences in read quality between the samples during genotyping, we did not use HERRO error correction for the 2015T reads for graph-based genotyping. As a final step of quality control, we manually assessed 32 instances in which the post-filtering graph-based genotypes deviated from the candidate variant callsets used to construct the graph by visual inspection in IGV, and removed 6 loci for which the graph genotype was likely incorrect or for which the true underlying genotype remained ambiguous (see Figure 5A for variant calls by sequence context before and after filtering). Graph-based genotyping recovered previously undetected SNPs (in one of the samples) and rescued filtered variants, improving the accuracy of genotype determination (see Figure S8).

**Figure 5.**
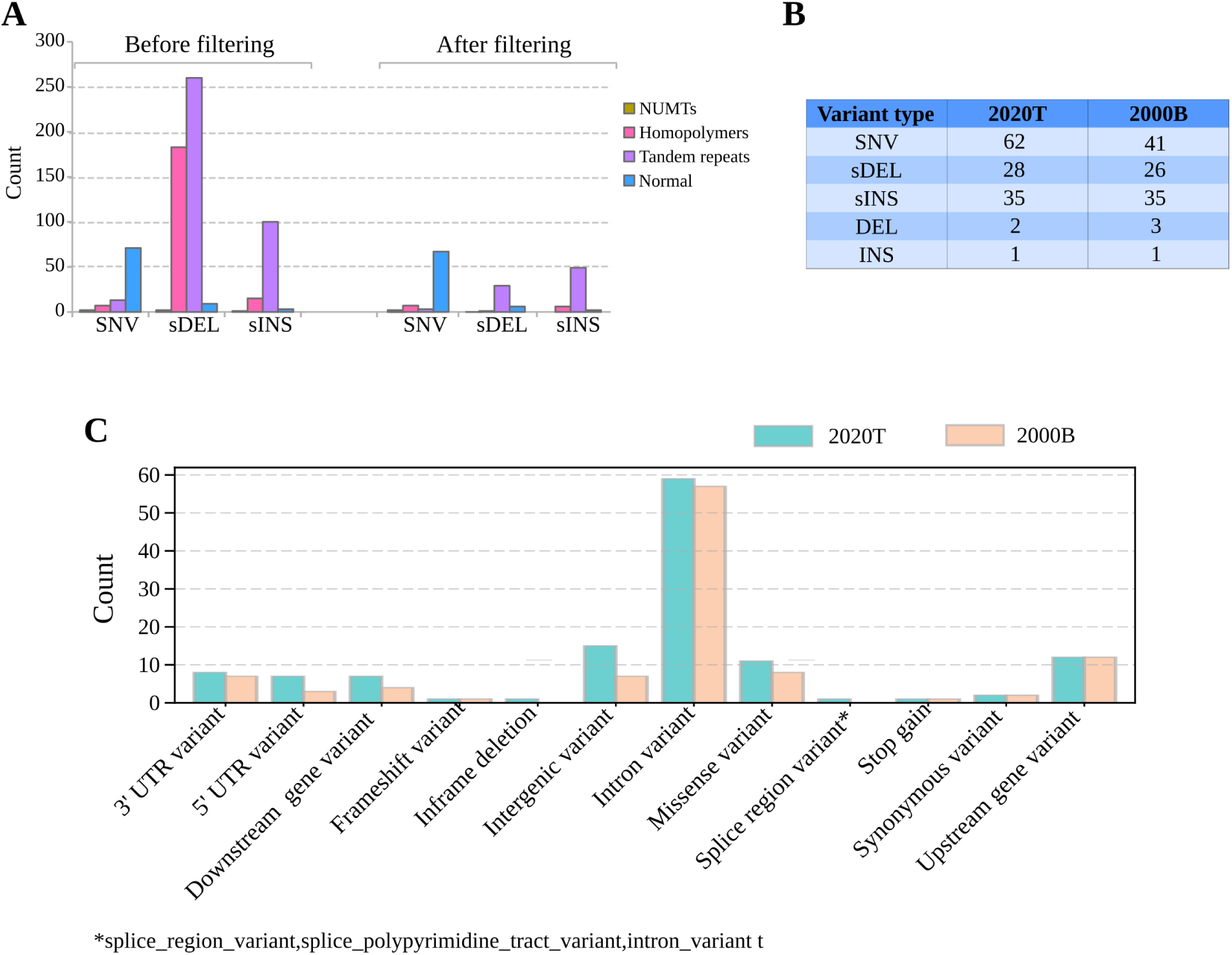
Summary of variant filtering and functional annotation. (A) Distribution of small variants by sequence context (NUMTs, homopolymers, tandem repeats, and normal regions) before and after filtering. Filtering removed spurious calls in repetitive regions, particularly among small deletions (sDEL). (B) Number of detected non-reference alleles in isolates 2000B and 2020T, stratified by variant type. (C) Functional annotation of small variants using Ensembl-VEP.

Our final callset included 177 loci that were genotyped for all 3 isolates; the callset was dominated by SNVs (n = 79), followed by small insertions (n = 57), and included a total of five structural variants (Supplementary Table 7). The concordance between the graph-based final callsets and the pre-graph candidate variant callsets was generally high; of note, however, the graph-based approach enabled the detection of 16 non-reference alleles not comprised in the candidate variant callset, including the detection of the large insertion in the ROP8-ROP2A region in isolate 2020T (which was also supported by manual investigation of 2020T reads aligned to the reference modified by the inclusion of the ROP8-ROP2A SV; data not shown). In total, we found 106 non-reference alleles in isolate 2000B and 128 non-reference alleles in isolate 2020T (Figure 5B); when assessing inter-isolate genetic distances based on SNVs, we found that isolates 2000B and 2015T exhibited the highest degree of relatedness (Figure S9).

### Functional annotation of small variants

We used Ensembl-VEP [56] to annotate the identified small variants in isolates 2020T and 2000B for potential functional impact. Almost 50% of small variants were located in introns in both isolates (60 of 128 variants for 2020T and 57 of 106 variants for 2000B), followed by intergenic variants in 2020T and variants classified as “upstream gene variant” in 2000B (Figure 5C). 10 variants in 2000B and 14 variants in 2020T were predicted to alter amino acid sequence; in almost all cases, the predicted changes were driven by missense variation, whereas gained stop codons only accounted for a total of 2 instances, 1 specific for each isolate. In 2000B, the stop-gained variant was found in the 6-phosphogluconate dehydrogenase protein; deletion mutants of this gene were found to be associated with severely attenuated virulence *in vivo* [57]. In 2020T, the detected stop-gained event was located in the AP2 domain transcription factor AP2X-8, a member of the ApiAP2 family. This family plays a critical role in the biology of *T. gondii*, particularly in regulating gene expression that governs life cycle transitions and developmental stages [58]. Of 18 non-synonymous small variants detected across both isolates, six were located in hypothetical proteins; other affected genes included e.g. Microneme-like protein and Conoid gliding protein (Table 1).

**Table 1:** Overview of non-synonymous small variants in isolates 2000B and 2020T.

| Contig | Position | Type | Gene | Predicted effect | 2020T | 2000B | Predicted gene product |
| --- | --- | --- | --- | --- | --- | --- | --- |
| Contig_11 | 4845879 | SNV | TgME49_XII0073200 | Missense variant | 1 | 0 | Hypothetical protein |
| Contig_11 | 1499544 | SNV | TgME49_XII0024800 | Missense variant | 1 | 1 | Hypothetical protein |
| Contig_20 | 185661 | SNV | TgME49_VI0008200 | Missense variant | 1 | 0 | Guanylate cyclase organizer (UGO) |
| Contig_20 | 2140215 | SNV | TgME49_VI0036200 | Stop gained | 0 | 1 | 6-Phosphogluconate dehydrogenase |
| Contig_20 | 3137120 | SNV | TgME49_VI0050600 | Missense variant | 1 | 0 | Microneme-like protein |
| Contig_20 | 1235798 | SNV | TgME49_VI0022100 | Missense variant | 1 | 1 | Conoid gliding protein (Cgp) |
| Contig_24 | 2197273 | SNV | TgME49_III0038200 | Missense variant | 1 | 1 | Cleft lip and palate transmembrane protein 1 (Clptm1) |
| Contig_30 | 2193723 | SNV | TgME49_IX0037400 | Missense variant | 1 | 0 | Heat repeat-containing Protein |
| Contig_30 | 491285 | INS | TgME49_IX0011000 | Frameshift variant | 0 | 1 | Hypothetical protein |
| Contig_42 | 4341951 | SNV | TgME49_VIIa0066400 | Missense variant | 0 | 1 | Phosphatidate cytidyltransferase |
| Contig_43 | 3772043 | SNV | TgME49_VIII0061200 | Missense variant | 1 | 1 | Hypothetical protein |
| Contig_43 | 3493540 | SNV | TgME49_VIII0057400 | Missense variant | 1 | 1 | Chromodomain helicase DNA binding protein Chd1/Swi2/Snf2 |
| Contig_43 | 8031232 | DEL | TgME49_VIII0121700 | Inframe deletion | 1 | 0 | Hypothetical protein |
| Contig_46 | 6422637 | SNV | TgME49_X0090900 | Missense variant | 1 | 0 | Hypothetical protein |
| Contig_46 | 5243975 | SNV | TgME49_X0073000 | Missense variant | 1 | 1 | Ulk kinase |
| Contig_46 | 1929612 | SNV | TgME49_X0029600 | Missense variant | 0 | 1 | PolyA polymerase |
| Contig_46 | 6778532 | SNV | TgME49_X0096700 | Stop gained | 1 | 0 | Ap2 domain transcription factor Ap2x-8 |
| Contig_46 | 1046388 | SNV | TgME49_X0016800 | Frameshift variant | 1 | 0 | Hydrolase, Nudix family protein |

### Functional annotation of structural variants

In addition to the large insertion in the ROP8-ROP2A region (see above), the graph callset included four additional deletions (Table 2), ranging in size from 52 to 1,180 bp. Three of these deletions were located in or affected exonic regions of predicted genes, including (i) a deletion specific to 2000B, occurring in the predicted overlapping hypothetical proteins TgME49_IV0036400.1 and TgME49_IV0036400.2 and leading to the loss of exon 4 in both cases; of note, the exons of TgME49_IV0036400.2 exhibited a complex repeat structure with a high degree of inter-exon homology (Figure S10); (ii) the likely inactivation of a hypothetical protein-encoding gene (TgME49_IX0117400), located within a cluster of orthologous genes predicted by Companion, due to the deletion of its single exon; and (iii) a deletion associated with a frameshift event in a “PT repeat”-annotated gene (TgME49_IX0109000); of note, this deletion was present in both 2020T and 2000B. The fourth detected deletion was located in an intronic region of a “histidine acid phosphatase superfamily”-annotated gene (TgME49_VIIa0014000) and was associated with the shortening of an AT-rich repeat region. In conclusion, in addition to differences at the level of small variants, the sequenced isolates also exhibited genetic differences at the structural level.

**Table 2:** Overview of structural variants in isolates 2000B and 2020T.

| Contig | Position | Type | SV length | Gene | Predicted effect | 2020T | 2000B | Predicted gene product |
| --- | --- | --- | --- | --- | --- | --- | --- | --- |
| Contig_46 | 7310767 | INS | 12896 | - | Expansion in ROP8-ROP2A locus | 1 | 1 | - |
| Contig_2 | 2235315 | DEL | 520 | TgME49_IV0036400 | Exon removal of a highly repetitive gene | 0 | 1 | Hypothetical protein |
| Contig_42 | 608620 | DEL | 52 | TgME49_VIIa0014000 | Shortening of an AT-rich repeat region in intron | 0 | 1 | Histidine acid phosphatase superfamily protein |
| Contig_35 | 1340294 | DEL | 1180 | TgME49_IX0117400 | Exon removal of a single exon gene | 1 | 0 | Hypothetical protein |
| Contig_35 | 699700 | DEL | 203 | TgME49_IX0109000 | Frameshift deletion | 1 | 1 | Pt repeat containing protein |

## Discussion

We used Oxford Nanopore long-read sequencing to generate a new *T. gondii* ME49 reference genome and to characterize genetic variation between three independently maintained *T. gondii* ME49 isolates. Our analyses demonstrated that our new reference genome exceeded other available *T. gondii* ME49 reference genomes in terms of contiguity and accuracy, as well as the existence of significant within-strain variation at the level of both large and small variants. Furthermore, functional annotation pointed to potential effects of the detected variants on virulence and life cycle transition phenotypes, suggesting a potentially confounding role in some of the experimental systems in which *T. gondii* ME49 is used as a model organism.

In our analysis, we identified both small and large genetic variants, including a notable copy number variation (CNV) in the ROP8-ROP2A locus, a potentially virulence-associated region in *T. gondii* [59,60]. Among the three sequenced isolates, 2015T harbored only three copies, whereas 2000B (based on manual construction of the locus) and 2020T (based on graph genotyping) contained six copies. A previous study showed that this locus is expanded in *T. gondii*, while it is entirely absent in its close relatives *Neospora caninum* and *Hammondia hammondi* [61]. Short-read sequencing data from reference *T. gondii* strains (GT1, ME49, VEG) indicated that these harbor six copies of ROP8/ROP2A genes [61], suggesting that this configuration is ancestral and that the reduced copy number in 2015T likely resulted from a deletion event. Additionally, we identified four large deletions, of which one was shared between isolates 2000B and 2020T, and 172 small variants, the majority of which were SNVs.

While we did not carry out any functional studies, a phenotypic impact of at least some of the detected variants seems plausible. First, the ROP8-ROP2A structural variant was associated with a difference in the total number of ROP8-ROP2A genes from three to six, with three of the additional gene copies being potentially expressed. Although a complete deletion of this locus in the RH strain did not show any apparent effects on tachyzoite growth *in vitro* [62], the ROP2 family (of which ROP8 is a member) is generally considered a key virulence factor [60,63,64]. ROP2A and ROP8 were found to interact with ROP18 and GRA7, two proteins that disrupt immunity-related GTPases and act synergistically to regulate acute virulence in mice [59]. Additionally, previous studies reported that ROP8 was upregulated during the tachyzoite-to-bradyzoite transition in the M4 strain [65], while ROP2A was upregulated in association with sulfadiazine resistance in type II strains [66]. While further functional studies are necessary to determine the impact of the ROP8-ROP2A structural variant, the difference in the combined ROP8-ROP2A gene dosage may plausibly affect *T. gondii* virulence and/or life cycle transitions. Second, additional structural variants were associated with the deletion of entire exons in genes *TgME49_IV0036400* and *TgME49_IX0117400* annotated as encoding hypothetical proteins, a predicted frameshift mutation in *TgME49_IX0109000* hypothetical protein, and the disruption of a tandem repeat stretch in an intron in the gene *TgME49_VIIa0014000* annotated as the histidine acid phosphatase superfamily protein. If the corresponding hypothetical proteins are expressed, a phenotypic effect of these variants seems likely. Notably, histidine acid phosphatase superfamily protein was predicted to play a role in host invasion based on ToxoNet, a high-confidence *T. gondii* protein-protein interaction map [67]. However, the functional impact of an intronic deletion in this gene remains unclear. Lastly, among the 18 non-synonymous variants we detected, several affect proteins involved in metabolic processes (e.g., 6-phosphogluconate dehydrogenase [57], phosphatidate cytidylyltransferase [68], NUDIX hydrolase [69], transcriptional regulation and chromatin remodeling (e.g., Ap2 domain transcription factor Ap2x-8 [58], Chromodomain Helicase DNA Binding Protein Chd1/Swi2/Snf2 [70], cellular signaling (e.g., Guanylate Cyclase Organizer [71], Ulk Kinase [72], and parasite motility (e.g. Conoid Gliding Protein [73]. These findings suggest that the detected variations could have functional consequences across multiple biological pathways, including metabolism, gene regulation, host-cell interaction, and intracellular signaling. Furthermore, six of the detected small non-synonymous variants were annotated as affecting genes encoding hypothetical proteins, highlighting that the substantial proportion of the *T. gondii* proteome that remains uncharacterized is a challenge for functional interpretation across variant classes.

While we unambiguously detected the presence of genetic variation, we did not attempt to determine when these variants arose in the propagation history of the sequenced isolates or whether their emergence was driven by neutral genetic drift or selection. In terms of selective pressure, the propagation environments of isolates 2015T and 2020T, which were maintained *in vitro* in cell culture, were much more similar to each other than to that of isolate 2000B, which was maintained *in vivo* under immune system pressure from the murine host. Additionally, 2015T and 2020T were acquired relatively recently from ATCC (the exact acquisition date for 2015T is unknown). Notably, however, pairwise genetic distances, measured by shared SNV alleles, were relatively similar across all isolates, with 2015T and 2000B unexpectedly exhibiting the highest pairwise genetic similarity. It is possible—though not testable within the scope of this study—that a proportion of isolate-distinguishing genetic variants were already present when the isolates were obtained from ATCC. In conclusion, it is possible that isolates 2015T and 2020T do not share a relatively recent common ancestor, and the evolutionary relationship between the characterized isolates remains unresolved.

Key methodological innovations of our study in the field of *T. gondii* genomics include the use of high-quality Oxford Nanopore R10 long-read sequencing and the utilization of a pangenomic approach for joint genotyping. With recent improvements in base calling and flow cell chemistry, Nanopore technology has achieved a raw-read accuracy >99% [74], and near-perfect genome assemblies have already been demonstrated for bacterial genomes [37]. In the context of our study, high-quality Oxford Nanopore sequencing, particularly with R10.4.1 chemistry and HERRO error correction, enabled the assembly of the most contiguous and complete ME49 genome to date; compared to the Xia et al. assembly, which was also based on Nanopore sequencing data but generated with older R9 flow cells, this was particularly evident from the increased proportion of perfectly aligned ToxoDB genes, indicating higher consensus accuracy. Joint graph-based genotyping was instrumental in the reliable detection of isolate-distinguishing genetic variation, including the identification of the ROP8-ROP2A structural variant in isolate 2020T, the rescue of individual variants that would have otherwise been removed (e.g., due to quality or coverage filters), and the detection of potential false-positive variant calls in isolates 2000B and 2020T by filtering for positions with a reference allele frequency of <90% in isolate 2015T. However, even with the most recent Nanopore flow cell chemistry, base calling algorithms, and sophisticated filtering approaches, distinguishing between true- and false-positive variants in homopolymeric and tandem repeat regions remained a challenge, due to an increased error rate of Nanopore sequencing in these regions [75]. While rigorous filtering was therefore essential to ensure the most reliable variant call, homopolymeric and tandem repeat regions are known to exhibit increased mutation rates in many species [76,77] and it is likely that our filtering approach led to the removal of some true-positive variants in these regions.

In addition to representing the highest-quality assembly for *T. gondii* strain ME49, our reference genome is also among the highest-quality *T. gondii* assemblies for any strain currently available [46,78]. Reference genomes play important roles in e.g. the analysis of RNA-seq data and short-read-based variant calling [35], and ME49 is often used as a reference for *T. gondii* type II strains [79,80]. The reference genome produced by us can therefore contribute to improved transcriptomic and population genetic characterization of *T. gondii*. However, some limitations remain. First, approximately 80 kb of sequence content from the Xia et al. assembly were not recovered in our assembly; further investigation is needed to determine whether this represents truly missing sequence or an assembly artifact in Xia et al. Second, due to its highly repetitive structure, the rDNA region remained challenging to assemble [81]. Spanning the complete locus would require, based on an estimate of 110 rDNA tandem array copies per *T. gondii* genome [82], a read of approximately 1 Mbp in size; while such read lengths can be achieved with Nanopore sequencing, they typically require further protocol optimization for obtaining ultra-long reads [83,84]. Based on the longest rDNA-containing read in our dataset, we could, however, confirm that *T. gondii* ME49 rDNA arrays follow the - generally less common - “L-type” conformation, in which the 5S gene is linked to (8S–5.8S–28S) rDNA array units [81]. In contrast to an earlier study [82], we found that the 5S gene was encoded on the opposite strand relative to the other rDNA genes. Furthermore, we found that the *T. gondii rDNA* repeat unit measured approximately 8.3 kb in size, consistent with reports for strains P and RH (GenBank accession X75453.1 and X75429.1, respectively), and detected an inversion in the rDNA repeat array, which may also exist in humans [85]. Last, our assembly did not contain a resolved mitochondrial genome; this may be due to the as-yet incompletely understood, and possibly fragmented, genomic architecture of mitochondrial DNA in *T. gondii* [86].

### Conclusion

In conclusion, we generated a new reference genome for *T. gondii* ME49 and demonstrated the existence of genetic divergence of potential phenotypic importance within the ME49 strain at the level of both large and small genetic variants. In addition to the specific findings for *T. gondii* ME49, our study highlights the importance of continuous monitoring of the genetic identity and stability of model organisms and shows how this aim can be achieved with modern high-accuracy long-read sequencing platforms.

## Methods

### Parasite cell culture

*T. gondii* ME49 isolates 2015T and 2020T were maintained by serial passaging of tachyzoites in human foreskin fibroblasts (HFFs) Hs27. HFFs were cultured in Iscove Modified Dulbecco Media (IMDM) containing 10% fetal bovin serum (FBS) at 37°C in a 5% CO2 incubator. The isolates were obtained from ATCC under the same catalog number (50611) (refer to the Supplementary Notes for provenance details about the isolates).

### *T. gondii* Cyst Preparation

*T. gondii* ME49 2000B cysts were isolated from the brains of CD1 mice via Ficoll-Paque gradient centrifugation 11 to 19 weeks post-infection. Briefly, mouse cerebrum tissue was homogenized by sequentially passing through progressively smaller cannulas (smallest gauge: 23G). The homogenate was first centrifuged at 130 x g for 5 min at room temperature (RT). The resulting pellet was resuspended in 20 ml of sterile PBS. To separate cysts, 10 ml of Ficoll-Paque Plus (GE Healthcare, USA) was carefully layered below the PBS suspension, followed by centrifugation at 1,250 x g for 25 min at room temperature, without using brakes. The cysts were then collected, washed in PBS, and stored at either 4°C (for short-term nucleic acid extraction) or −80°C (for long-term nucleic acid extraction) until further processing.

### HMW DNA Extraction

*T. gondii* ME49 tachyzoites (2015T and 2020T) were cultured in a confluent HFF monolayer for approximately 7 days, until the monolayer was fully infected and the tachyzoites had lysed the host cells to emerge. The medium containing the released tachyzoites was collected for DNA extraction. For 2000B, DNA was extracted from the collected cysts. Extraction for isolate 2020T and 2000B was done using Phenol-Chloroform Extraction (see below). In contrast, extraction for the 2015T isolate was performed using one of the following methods in an effort to obtain high-quality, non-fragmented DNA for genome assembly (Supplementary Table 1).

1. **Monarch HMW DNA Extraction Kit** (NEB, Cat# T3050L): DNA was extracted using the Monarch HMW DNA Extraction Kit according to the manufacturer’s protocol.
2. **Size Selection via Semi-Selective DNA Precipitation after extraction:** DNA was size-selected to deplete short fragments using a semi-selective DNA precipitation method, following the August 2021 version of the protocol from Oxford Nanopore Technologies [87]
3. **Ultra-Long DNA Sequencing: For ultra-long DNA sequencing:** The Ultra-Long DNA Sequencing Kit (SQK-ULK114) was used according to the November 2022 ONT protocol [84].

### Phenol-Chloroform extraction for gDNA 2020T and 2000B samples

Genomic DNA was purified using a phenol-chloroform extraction protocol. Briefly, *T. gondii* cells (5 × 10^7) were washed in PBS and resuspended in a digestive solution containing 500 µl TNE, 50 µl 10% SDS, and 7.5 µl Proteinase K (10 mg/ml). The suspension was incubated overnight at 56°C with shaking. Following lysis, 500 µl of chloroform/isoamyl alcohol (24:1) was added, mixed vigorously and centrifuged at 13,000 rpm for 5 minutes at room temperature. The supernatant was transferred to a new tube, and the extraction was repeated. DNA was precipitated with 500 µl isopropanol, incubated for 5 minutes at room temperature, and centrifuged at 8,000 rpm for 10 minutes. The DNA pellet was washed three times with 500 µl of 70% ethanol, dried for 10 minutes at room temperature, and resuspended in 100 µl TE buffer (incubated for 1 hour at 56°C, then overnight at 4°C).

### Oxford nanopore sequencing

Data acquisition for the three different *T. gondii* isolates was performed using Oxford Nanopore sequencing (R10.3, R10.4 or R10.4.1) from multiple samples with the aim of achieving at least 20X coverage. For 2015T, base-calling was performed using Dorado v0.7.3 [40] with the command dorado duplex. Error correction on simplex reads was then carried out using HERRO [41] independently for the data generated by each sequencing run. After correction, simplex and duplex reads were combined. For 2020T and 2000B, all sequencing runs were basecalled using Guppy v6.1.5 [42] For details about the sequencing and basecalling of each sample, see Supplementary Table 1.

### *Toxoplasma gondii* ME49 genome assembly

To eliminate human DNA contamination, the sequencing data were filtered by aligning the reads against a combined *T. gondii* ME49 (GenBank: JACEHA000000000.1) (Xia et al., 2021), human (GRCh38 Primary Assembly, RefSeq: GCF_000001405.26) and mouse (GRCm39 Genome Assembly, RefSeq: GCF_000001635.27) reference genomes using minimap2 v2.28-r1209 [88]. Reads mapping to the human or mouse genomes were removed, and the remaining reads were extracted from the FASTQ file using Seqkit v0.15.0 [89] with the command seqkit grep -v. The filtered FASTQ files from all 2015T sequencing runs were combined using the cat command and further filtered to remove mitochondrial reads. This filtering was performed by aligning the reads against a FASTA file containing mitochondrial sequence blocks and mitochondrial cytochrome genes [86]. Reads with 40% or more of their length covered by mitochondrial sequence blocks were removed from the FASTQ file using a custom Python script. The remaining sequencing reads were then used to assemble the genome with Flye v2.9.4-b1799 [43] using the --nano-corr parameter. One circular contig likely associated with contaminant DNA from the bacterial species *Facklamia ignava* was removed.

### Gene annotation of the 2015T assembly

To annotate the new assembly, the Companion web tool v2.2.0 was employed [45], with default settings and utilizing the *T. gondii* ME49 reference available in the Companion database. The output GFF3 file was then modified to ensure compatibility with subsequent gene annotation tasks, using gffread with the options -O -F -E --force-exons. Transcript biotype information, including rRNA, tRNA, pseudogenic transcript, and protein-coding (mRNA), was incorporated into the GFF3 file using a custom script. Finally, the GFF3 file was compressed using bgzip and indexed with Tabix.

### Comparison of the 2015T assembly to the Xia et al. Assembly

To compare the 2015T assembly with the Xia et al. assembly, we aligned the two genomes using nucmer from the MUMmer4 package [47] with default parameters, followed by dnadiff. Chromosome mapping between assemblies was performed using minimap2 with parameters -x asm20 -m 10000 -z 10000,50 -r 50000 --end-bonus=100 --secondary=no -a -t 20 -- eqx -Y -O 5,56 -E 4,1 -B 5. Dot plots were generated using D-Genesis [90] for both nuclear and apicoplast sequences to visualize large-scale structural differences. To obtain reference gene sequences for *T. gondii* from ToxoDB v48, gene coordinates were extracted from the ToxoDB- 68_TgondiiME49.gff annotation file and converted to BED format. The corresponding sequences were retrieved from the ToxoDB-68_TgondiiME49_Genome fasta file using bedtools getfasta. The final dataset contained 8,778 gene sequences. Gene sequences were then mapped to the assembly using minimap2 map-ont, and a custom Python script was used to count the number of mapped genes and calculate the fraction of perfectly mapped genes. Additionally, assembly anomalies were identified by mapping the reads back to the assembly and inspecting the alignments in IGV v2.17.4 [49] (see Supplementary Note: Identification of Assembly Anomalies).

### Analysis of the rDNA region

The breakpoint in the 2015T assembly occurred in Chromosome IX, within a repetitive rDNA region that was misassembled in the Xia et al. assembly. The longest read from our dataset that mapped to this position (JACEHA010000011.1:1376356-1526114) in the Xia et al. assembly was 207,813 bp in length and spanned positions 1,387,207 to 1,515,512. The read was extracted using seqkit and aligned, using minimap2 map-ont, to the longest available rRNA sequences for the 28S, 5.8S, and 18S genes of *T. gondii* ME49 (NCBI: XR_001974492.1, XR_001974334.1, and XR_001974441.1), as well as to the 5S rRNA sequence from the RH strain (GenBank: 5XXB_3). The alignment coordinates of these sequences on the longest read were visualized using a custom Python script. Similarly, the sequence from the Xia et al. assembly at position (JACEHA010000011.1:1276356-1626114) was extracted and aligned to the rRNA sequences, with the alignment coordinates visualized using the custom Python script.

### Calling of SNPs and INDELs

2020T and 2000B isolate FASTQ files were filtered to remove contamination from human or mouse sources, employing the same approach that was also used for the 2015T reads. Single nucleotide polymorphisms (SNPs) and insertions/deletions (INDELs) in 2000B and 2020T were called separately for each sample using Medaka (v2.0.1) [51] with the medaka_variant script and using the 2015T assembly as a reference genome. Variant calling was carried out using the Medaka model r104_e81_sup_variant_g610, which was empirically found to reduce the number of false-positive variant calls in isolate 2000B compared to more recent Medaka models. Variants were annotated and classified using the varType command of SnpSift v5.1 [91]. Only variants located on nuclear chromosomes were considered. We applied a stringent filtering approach to Medaka-identified variants based on sequencing depth, genotype quality, and deletion overlap. Variants were retained if the sequencing depth (DP) was ≥10 and the genotype quality (GQ) was ≥10. Additionally, for SNPs, we assessed the proportion of deletions at the variant position; SNPs at positions where deletions accounted for ≥50% of the total depth were removed. Additionally, we removed small variants that overlapped with structural variants (SVs), occurred at the very end or beginning of the chromosome where assembly quality is potentially suboptimal, or that were located at previously identified erroneous or suspicious positions.

### Calling SVs

For each sample, Sniffles v2.2 [52] was used to identify structural variants. To ensure the VCF file conformed to the VCF version 4.1 specifications [92], a custom a custom script was developed to adjust SV calls by decrementing the variant position and updating the reference and alternate alleles for insertions and deletions, accordingly. All called SVs were then manually evaluated in IGV v2.17.4. Variants were retained if they were clearly supported by long reads spanning the entire variant region, such that the structural change was directly observable in the read alignments or supported by consistent and extensive read clipping. SVs were excluded if they were located on the apicoplast contig included from the Xia et al. assembly, corresponded to translocations resulting from incomplete chromosome assembly, or lacked strong read support. In addition, SVs were also excluded if they overlapped regions potentially enriched for assembly artifacts—such as contig ends—or if they fell within nuclear mitochondrial DNA segments (NUMT) regions, in which misaligned mitochondrial-origin reads commonly produced spurious signals. Filtering results for all SV are summarized in Supplementary Table (Sniffles manual filtering).

### ROP8-ROP2A locus annotation

Our objective was to characterize the number and order of ROP2A and ROP8 gene copies within the RO8-ROP2A locus of three assemblies: Xia et al. assembly, our 2015T assembly, and the sequence of the locus in isolate 2000B. Inspecting Companion-based gene annotations for this locus, we found that Companion classified all ROP2A sequences as pseudogenes, likely driven by the incompleteness of the ROP2A reference sequence obtained from ToxoDB. We thus employed BLASTn and Augustus [53] in addition to Companion for the characterization of the ROP8-ROP2A. We first used BLASTn v2.14.1+ to obtain candidate ROP8 and ROP2A sequences from the assemblies. We mapped the reference ROP2A (*TGME49_215785*) and ROP8 (*TGME49_215775*) sequences from ToxoDB (release 68) to the assemblies, extracted the sequence coordinates of the BLAST hits, and merged the coordinates of ROP8 and ROP2A hits whenever they overlapped; finally, we extracted the sequences corresponding to the merged coordinate sets from the assemblies. In addition, we ran Augustus (using -- species=toxoplasma --strand=both --genemodel=complete --codingseq=on) on the assembly sequences and extracted the sequences of the detected genes. To create a merged set of candidate gene sequences, we combined the BLAST- and Augustus-based sets, merging BLAST- and Augustus-derived sequences whenever they overlapped on the assembly they were extracted from. In a final step, we cross-checked the merged set against Companion to ensure completeness; no additional sequences were added during this step.

To determine sequence homology between the extracted candidate gene sequences, and to determine homology between the candidate gene sequences and ROP2A and ROP8 reference sequences, we computed a multiple sequence alignment (MSA) using Clustal Omega v1.2.4 [93], including i) the candidate gene sequences from the investigated assemblies; ii) the ToxoDB reference sequences for ROP2A and ROP8; and iii) the Augustus-predicted gene sequences from the investigated assemblies (i.e., Augustus-based sequences prior to merging with BLAST hits), as these were informative about the positions of start and stop codons within the extracted sequences. We projected start and stop codon positions from Augustus- and ToxoDB-derived sequences into the MSA and obtained MSA interval (positions 409 – 2122) corresponding to the union of the implied open reading frames; pairwise sequence similarities between the extracted candidate gene sequences were computed based on this MSA interval. In addition, we projected the start and stop positions of ROP2A and ROP8 ToxoDB sequences into the MSA and intersected the implied MSA coordinates (positions 409 – 1700); assignment of the candidate gene sequences to either ROP2A or ROP8 was based on pairwise sequence similarity between the individual candidate gene sequences and ROP2A / ROP8 ToxoDB reference sequences over this interval. . Of note, the end of the latter range (MSA position 1700) corresponded to the end of the partial ROP2A sequence, which is not complete.

Similarity matrices and heatmaps were generated using a custom Python script. To assess which of the identified candidate gene sequences were potentially functional, we mapped Toxoplasma ME49 RNA-seq sequencing data retrieved from the NCBI Sequence Read Archive (SRA) using prefetch (SRX6428507) and converted them to FASTQ format with fasterq-dump. Paired-end reads from SRR9667927 and SRR9667928 were merged to create a combined dataset. The merged reads were then mapped to the stitched 2000B sequence using STAR v2.7.10a [94] with the following parameters: --alignIntronMax 10000 and --alignMatesGapMax 1000 --outMultimapperOrder Random.

### Graph construction and genotyping from the graph

A genome graph was constructed using the vg toolkit v1.63.1 (75) based on a catalog of previously identified SNPs, indels, and structural variants (SVs). Variant calls from Medaka and Sniffles2 were merged using bcftools concat with duplicate removal, followed by position-wise merging across samples using a modified version of the merge_vcfs.py script from PanGenie [95]. The script was adapted to process a haploid genome and handle genotype formats from Sniffles2 and Medaka . The final variant catalog was compressed and indexed with bgzip and tabix. The genome graph was built with vg construct, retaining all alternative paths (-a), and indexed (-L) in xg-format (-x). Long-read sequencing data were aligned to the graph using GraphAligner v1.0.18-[55] employing an identity threshold of 0.75 (--precise clipping 0.75). Alignments were filtered for a minimum mapping quality of 30 (vg filter -q 30), and variant genotyping was performed with vg pack (-Q 20), followed by vg call (-d 1).

### Identification of Low-Complexity Genomic Regions and NUMTs

Tandem repeats in the 2015T assembly FASTA file were identified using GMATA (v2.3) [96] with the following parameters: motif unit lengths ranging from 2 to 6 nucleotides and a minimum of 5 repeat units. Homopolymers were identified using the homopolymer_finder.py script from the umiVar toolkit [97] with a minimum homopolymer length of 4 bases (-l4). The resulting positions were converted into BED format using an AWK script, which grouped consecutive positions into continuous intervals. Positions on apicoplast sequences were excluded from further analysis. NUMTs were detected using RepeatMasker (v4.1.7-p1) [98] with a custom library of mitochondrial sequences [86]. Positions detected on apicoplast sequences, as well as those corresponding to low-complexity regions identified by RepeatMasker’s default settings, were excluded. The remaining NUMT coordinates were saved in BED format.

### Variant annotation using Ensembl Variant Effect Predictor (VEP)

Variants were functionally annotated using Ensembl VEP [56] with the 2015T assembly as the reference genome. The annotation was performed using the Companion-generated GFF3 assembly annotation file generated and the --pick parameter to retain a single, most relevant consequence per variant.

## Declarations

### Ethics approval and consent to participate

Not applicable

### Consent for publication

Not applicable

### Availability of data and materials

The raw Oxford Nanopore sequencing data generated and analyzed during this study have been deposited in the NCBI Sequence Read Archive (SRA) under BioProject accession number PRJNA1241696. The de novo genome assembly of the *Toxoplasma gondii* ME49 isolate 2015T has been deposited in GenBank under accession number JBMUND000000000. All custom scripts and computational workflows used in this study are openly available. The genome filtering and assembly pipeline is accessible at https://github.com/YomnaGohar/T.-gondii_filter_assemble, and the variant calling and graph-based genotyping pipeline is available at https://github.com/YomnaGohar/ToxoVar. Detailed instructions for reproducing the main figures are provided at https://github.com/YomnaGohar/Intra-Strain-Genetic-Heterogeneity-in-Toxoplasma-gondii-ME49.

### Authors’ contributions

YG performed the genome assembly, developed the analysis strategy. YG and AD wrote the manuscript. YG and MN jointly developed the variant analysis and graph-based genotyping pipeline. LH, DW, JM, KB, and MH prepared parasite samples and generated sequencing data. US, DD, and KP contributed to project discussions and provided feedback on the experimental design and interpretation of results. AD supervised the project and provided critical input throughout. All authors read and approved the final manuscript.

## Acknowledgements

Not applicable

## Figures, tables and additional files

**Supplementary Figure S1:**

Flowchart of the Snakemake pipeline used in the variant analysis. Created in https://BioRender.com

**Supplementary Figure S2: Improved alignment of sequencing reads and ToxoDB reference gene sequence in the 2015T assembly compared to the Xia et al. assembly.** In each IGV panel, the top track shows the gene from ToxoDB, and the bottom track displays the long reads used to construct each assembly.

(A) Read alignment over the *THH1 domain-containing* gene *(TGME49_222310)*. The Xia et al. assembly exhibits a high density of mismatches and indels, whereas the 2015T assembly shows clean and consistent alignment. (B) Read alignment over the *Malate dehydrogenase (MDH)* gene *(TGME49_318430)*. In the Xia et al. assembly, extensive mismatches and gaps lead to incomplete mapping of the gene sequence. In contrast, the 2015T assembly enables smooth and concordant alignments, reflecting improved sequence accuracy.

**Supplementary Figure S3. Evidence of a likely misassembled rDNA repeat region in the Xia et al. assembly.**

(A) IGV screenshot showing the alignment of reads used to construct the Xia et al. assembly (JACEHA010000011.1:137,482–1,530,609). A marked increase in read depth is observed in this region, accompanied by a lack of spanning reads, suggesting a potential misassembly. The bottom track shows the alignment of 2015T assembly contigs (contig_35 and contig_30). (B) Schematic representation of rDNA gene blocks annotated in the same region of the Xia et al. assembly.

**Supplementary Figure S4. Dot plot comparison of the apicoplast genome between the Xia et al. assembly and the 2015T assembly.**

A dot plot generated using D-Genies shows alignment between the apicoplast contig from the Xia et al. assembly (JACEHA010000016.1) and the 2015T assembly. A single contig from the 2015T assembly (contig_1) aligns to the Xia et al. contig in two segments: once in the forward direction and once in the reverse complement. This pattern is consistent with the presence of an inverted repeat in apicoplast genome.

**Supplementary Figure S5. Resolution of a structural variant in the 2000B isolate through reference modification.**

(A) IGV snapshot showing 2000B reads aligned to the original 2015T assembly, displaying a region with abnormal coverage and extensive read clipping, consistent with a misrepresented structural variant. (B) Alignment of the same 2000B reads to the modified reference shows a uniform coverage profile with minimal clipping, indicating that the structural variant was correctly resolved through manual reference correction. For clean visualization, secondary alignments and reads with mapping quality 0 (MAPQ0) were filtered out, and the coverage allele-fraction threshold was set to 0.5.

**Supplementary Figure S6. Evidence for a structural variant in 2020T relative to the 2015T reference.**

IGV screenshots showing read alignments of 2015T (top panel) and 2020T (bottom panel) to the 2015T assembly. While the 2015T reads align cleanly with uniform coverage, the 2020T reads show a mixed base pattern, where subsets of reads consistently support different genotypes across the same region. This pattern is similar to that observed in the 2000B alignment (Figure S5) and is consistent with a collapsed repeat or unresolved structural variation in the reference assembly. For clean visualization, secondary alignments and reads with mapping quality 0 (MAPQ0) were filtered out, and the coverage allele-fraction threshold was set to 0.5.

**Supplementary Figure S7. RNA-seq read coverage across the ROP2A–ROP8 locus in the 2000B.**

IGV screenshot showing RNA-seq alignments from *T. gondii* ME49 mapped to the structurally resolved ROP2A–ROP8 region of isolate 2000B. The coverage profile indicates detectable expression of ROP8-1, ROP8-2, ROP8-3, ROP2A-A, ROP2A-2B and ROP2A-1. The alignment is visualized with secondary alignments filtered out.

**Supplementary Figure S8. Examples of genotype correction through graph-based variant calling.**

(A) An example of a variant that was initially called only in isolate 2020T but missed in 2000B, despite clear evidence of its presence in both isolates. The graph-based genotyping approach correctly recovered the variant in both samples. (B) A case where a true variant in 2020T was erroneously filtered out during the Medaka-based filtering step. The variant was subsequently rescued by the graph-based genotyper, restoring the correct genotype.

**Supplementary Figure S9. Pairwise genetic similarity between isolates based on SNVs.**

Matrix showing inter-isolate genetic similarity calculated from shared single-nucleotide variants (SNVs). The highest level of similarity was observed between isolates 2000B and 2015T, indicating a closer genetic relationship relative to the other isolate pairs.

**Supplementary Figure S10. Pairwise exon similarity matrix for *TgME49_IV0036400.2***

Heatmap showing the percent identity between all exon pairs of the gene *TgME49_IV0036400.2*. Exons were aligned in all-vs-all fashion, and similarity was calculated as percent identity over alignment length. The exon indices on both axes start at 0 and follow the exon order in the GTF annotation.

**Supplementary Table 1. Summary of sequencing metadata and read classification for *T. gondii* ME49 isolates.**

**Supplementary Table 2. Genomic positions of anomalous regions identified in the Xia et al. assembly and their corresponding locations in the 2015T genome.**

**Supplementary Table 3. Medaka variant calls for 2020T and 2000B isolates before and after filtering**

**Supplementary Table 4. Classification of filtered Medaka variant calls by genomic context: NUMTs, homopolymers, tandem repeats, and normal positions**

**Supplementary Table 5. Structural variant calls before and after filtering.**

**Supplementary Table 6. Manual curation of structural variant calls with filtering decisions and supporting reasons.**

**Supplementary Table 7. Genotype concordance before and after graph-based genotyping, stratified by variant type.**

