## Supplementary Notes for "Intra-Strain Genetic Heterogeneity in *Toxoplasma gondii* ME49: Oxford Nanopore Long-Read Sequencing Reveals Copy Number Variation in the ROP8-ROP2A Locus"

**Propagation history of the sequenced *T. gondii* ME49 isolates**

2015T was obtained from ATCC (the exact date is unkown) and thawed in 2015. After thawing, it was cultured for two passages and then cryopreserved. In January 2022, it was thawed again and maintained in continuous cell culture, with passaging occurring twice per week. Samples for sequencing were collected between June and November 2024. Exact collection dates are provided in Supplementary Table 1. 2020T was obtained from ATCC in April 2020, thawed and cultured in May 2020, and has been continuously maintained in cell culture since that time, with biweekly passages. Two samples were collected for sequencing in December 2021 and May 2022 (Supplementary Table 1). 2000B was acquired in the early 1990s; the exact source is unknown. The isolate has been continuously propagated in mice at the Institute of Medical Microbiology and Hospital Hygiene, University Hospital Düsseldorf, for at least 30 years. Reinfection typically occurred every 3–6 months, depending on experimental needs. Periods of frequent passaging were associated with higher animal usage, while lower demand resulted in less frequent propagation. Sample collection for sequencing happend between January 2022 until July 2023. Exact collection dates are provided in Supplementary Table 1.

**Mitochondrial reads confound assembly at NUMT-Containing Locus**

During an initial genome assembly attempt that did not exclude mitochondrial reads, we identified an anomalous region that aligned with one of the identified anomalous loci in the Xia et al. ME49 assembly (see later). This artifact arises due to the presence of a large nuclear mitochondrial DNA segment (NUMT), which causes mitochondrial reads to erroneously align to the nuclear genome and induces a misassembly. This issue is clearly visible in the IGV screenshots in the following Figure: (A) shows the region before mitochondrial read filtering, where abnormal read alignment and coverage patterns are evident, while (B) shows the same region after filtering for mitochondrial reads and reassembly, where the assembly anomaly is resolved.

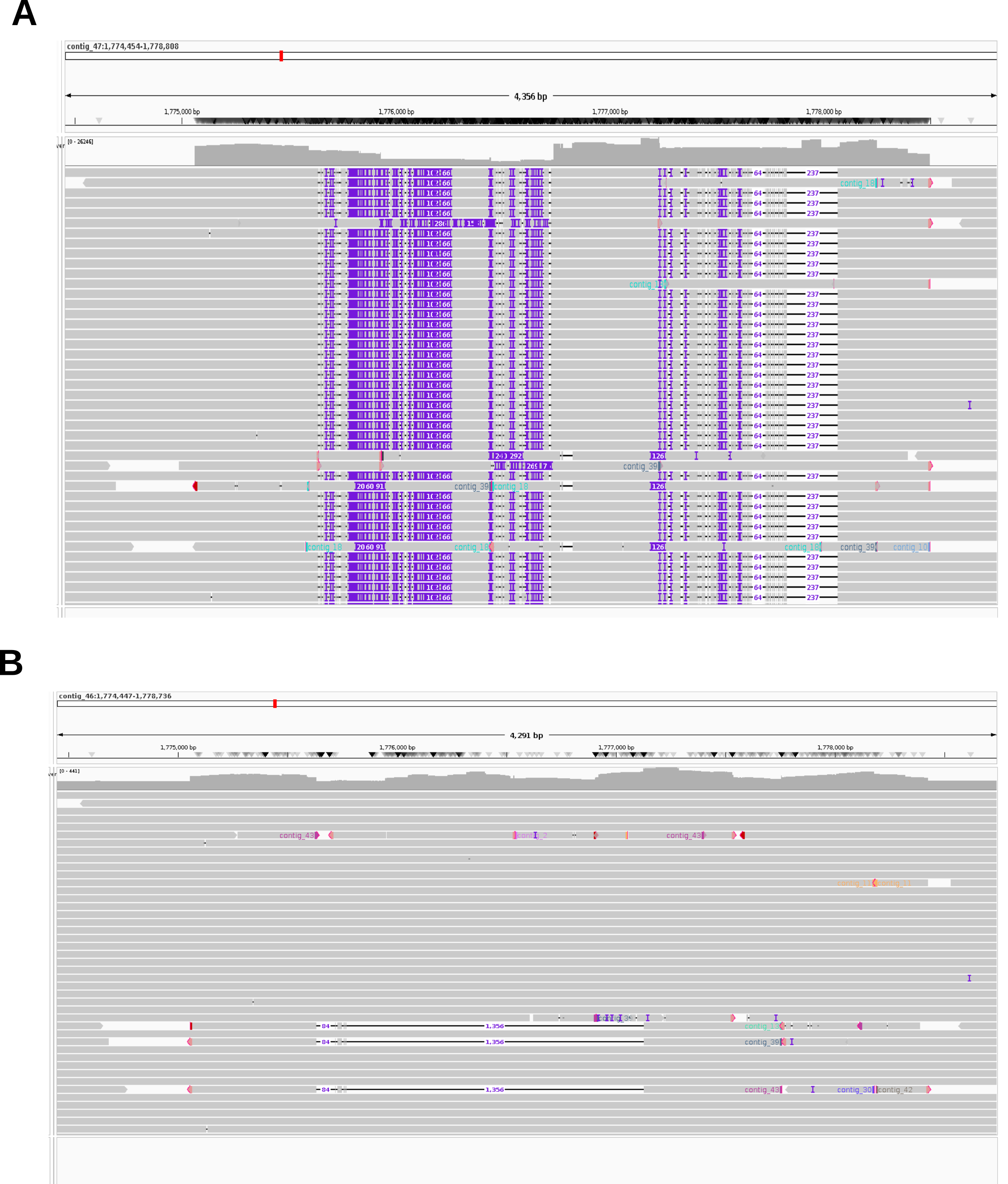

**Identification of Assembly Anomalies**

The entire Xia et al. assembly was, based on the Nanopore sequencing data generated by Xia et al. that was used to generate the assembly (SRA: SRX8862416), manually inspected in IGV v2.17.4 to assess assembly quality. The goal was to systematically identify anomalous positions that could indicate potential errors in the assembly. Anomalous positions were identified based on three primary criteria:

1. Read depth anomalies: Sudden increases or decreases in read depth, particularly regions with sharp depth drops or unexpected coverage spikes.
2. Evidence for structural variants (SVs) : Positions showing split-read signatures, or large insertions/deletions indicative of structural variations.
3. Patterns associated with collapsed repeats: Regions with stretches of positions containing multiple alleles, suggesting possible misassemblies due to repeat compression.

Positions were then classified into two categories:

- Erroneous positions: Positions where all reads mapping to the site exhibited the same anomaly, such as a uniform pattern of insertions or a complete lack of spanning reads at a high-depth site.
- Suspicious positions: Positions where some but not all reads exhibited anomalies, indicating possible inconsistencies or mixed signals in the sequencing data (e.g., at least one spanning read was present)

These positions were extracted with coordinates selected to cover the full extent of the anomalous regions. After creating the 2015T assembly, we used LiftOver [1](version 469, installed from Bioconda) to project these identified positions from the Xia et al. assembly onto the 2015T assembly. To facilitate coordinate projection, we implemented a Snakemake v7.32.4 pipeline, first mapping the new assembly to the JACEHA000000000.1 reference with minimap2 using the -cx asm20 -m 10000 -z 10000,50 -r 50000 --end-bonus=100 --secondary=no --eqx -Y -O 5,56 -E 4,1 -B 5 preset to generate a PAF alignment file. This file was converted to chain files using paf2chain [2] for use in coordinate conversion, and LiftOver was subsequently performed with the UCSC liftOver tool with -minMatch=0.4. Each position in the 2015T assembly was then reviewed in IGV to verify alignment consistency and read depth, ensuring that no unexpected variations or mismatches were present. A table containing the coordinates of the anomalous positions on the Xia et al. assembly and their corresponding positions on the 2015T assembly is provided in Supplementary Table 2. A final scan of the 2015T assembly in IGV confirmed the absence of new anomalies and validated that all regions exhibited consistent read alignment and expected depth, indicating a more accurate and homogeneous assembly.

**Manual curation of ROP8-ROP2A sequence in 2000B sample**

We defined two anchor genes—genes that are neither ROP2A nor ROP8 and that exist uniquely in the genome. The identified anchors were TGME49_215750, located left of the ROP2A-ROP8 locus, and TGME49_275310, located right of the locus. No single read from sample 2000B spanned both anchor genes in the 2015T assembly, i.e. no single read was informative about the true structure of this locus in the 2000B sample. To reconstruct the sequence of the locus in isolate 2000B, we identified a pair (A, B) of overlapping reads, selected according to the following criteria: i) read A contained the left anchor; ii) read B contained the right anchor; iii) A and B overlapped in the reference genome with similar mismatch pattern at the overlapping region. To find suitable read pairs, reads mapping upstream of position 7,302,329 in 2015T (left anchor) and those spanning position 7,302,329 to 7,322,405 in 2015T (right anchor) were extracted using SAMtools, and a pair of reads, meeting the criteria stated above, was selected: 9350f71a-a210-444c-97d9-ab2c695af5e3 (left) and cb2b7796-e42a-46b4-b204-7b3e511adecc (right) (panel A in the following figure). The second read was reverse-complemented, and the corrected sequence was used in analysis. The two reads were stitched together using Minimap2 (-cx ava-ont), selecting the alignment from the PAF file based on the expected 10,000 bp overlap. Bases up to position 17,115 in 9350f71a-a210-444c-97d9-ab2c695af5e3 were concatenated with cb2b7796-e42a-46b4-b204-7b3e511adecc starting from position 108. Reads from the 2000B sample mapping to the ROP8-ROP2A locus in the 2015T assembly were extracted and aligned to the stitched sequence using Minimap2 (-ax map-ont) (panel B in the following figure). The stitched sequence was then polished in two rounds with Medaka (r104_e81_sup_variant_g610_model_pt.tar.gz). Secondary alignments and small indels (<10bp) were filtered during visualization(panel C in the following figure).

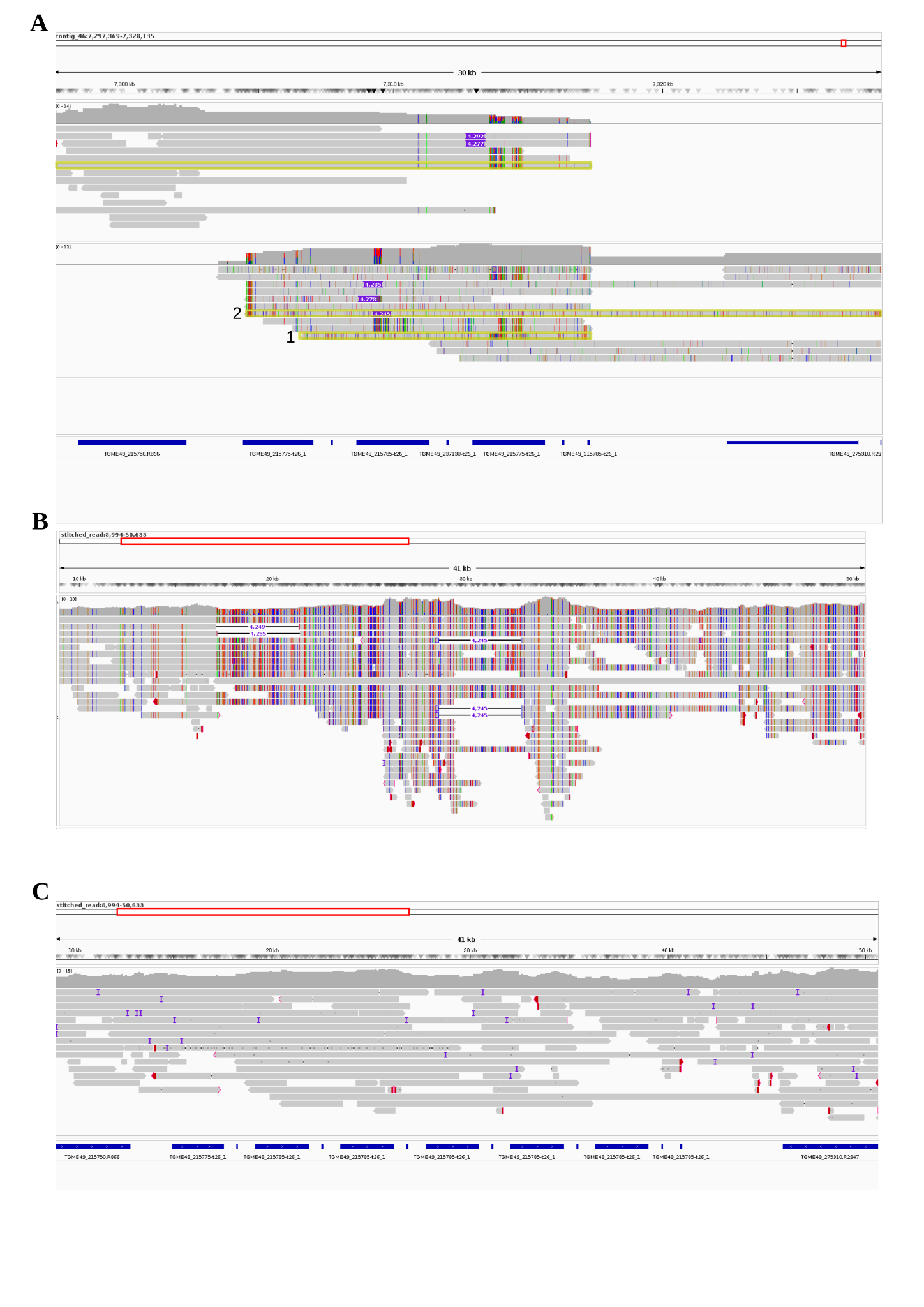
