## Supplementary figures and images for "Intra-Strain Genetic Heterogeneity in *Toxoplasma gondii* ME49: Oxford Nanopore Long-Read Sequencing Reveals Copy Number Variation in the ROP8-ROP2A Locus"

### Supplementary Figure 1

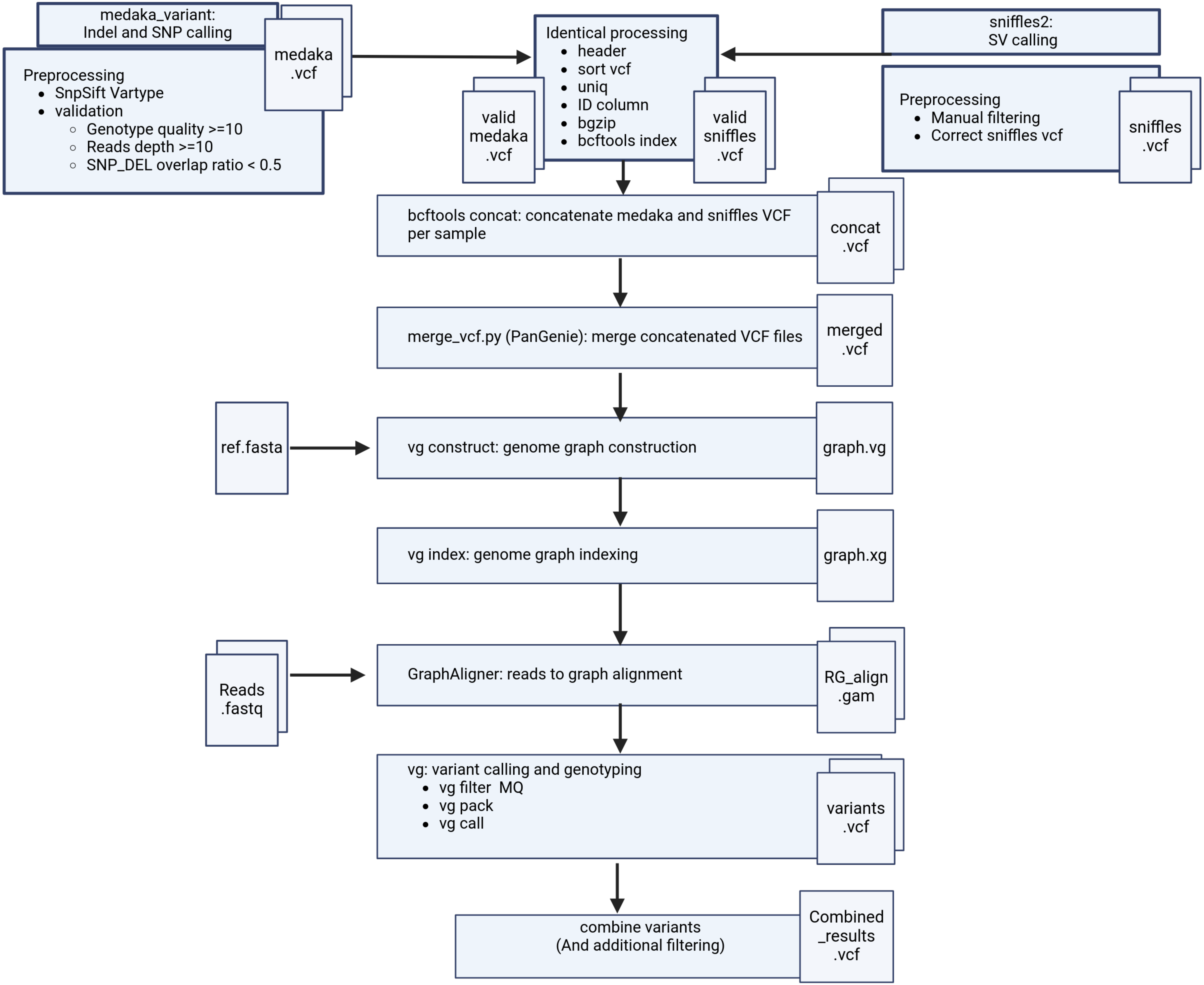

### Supplementary Figure 3

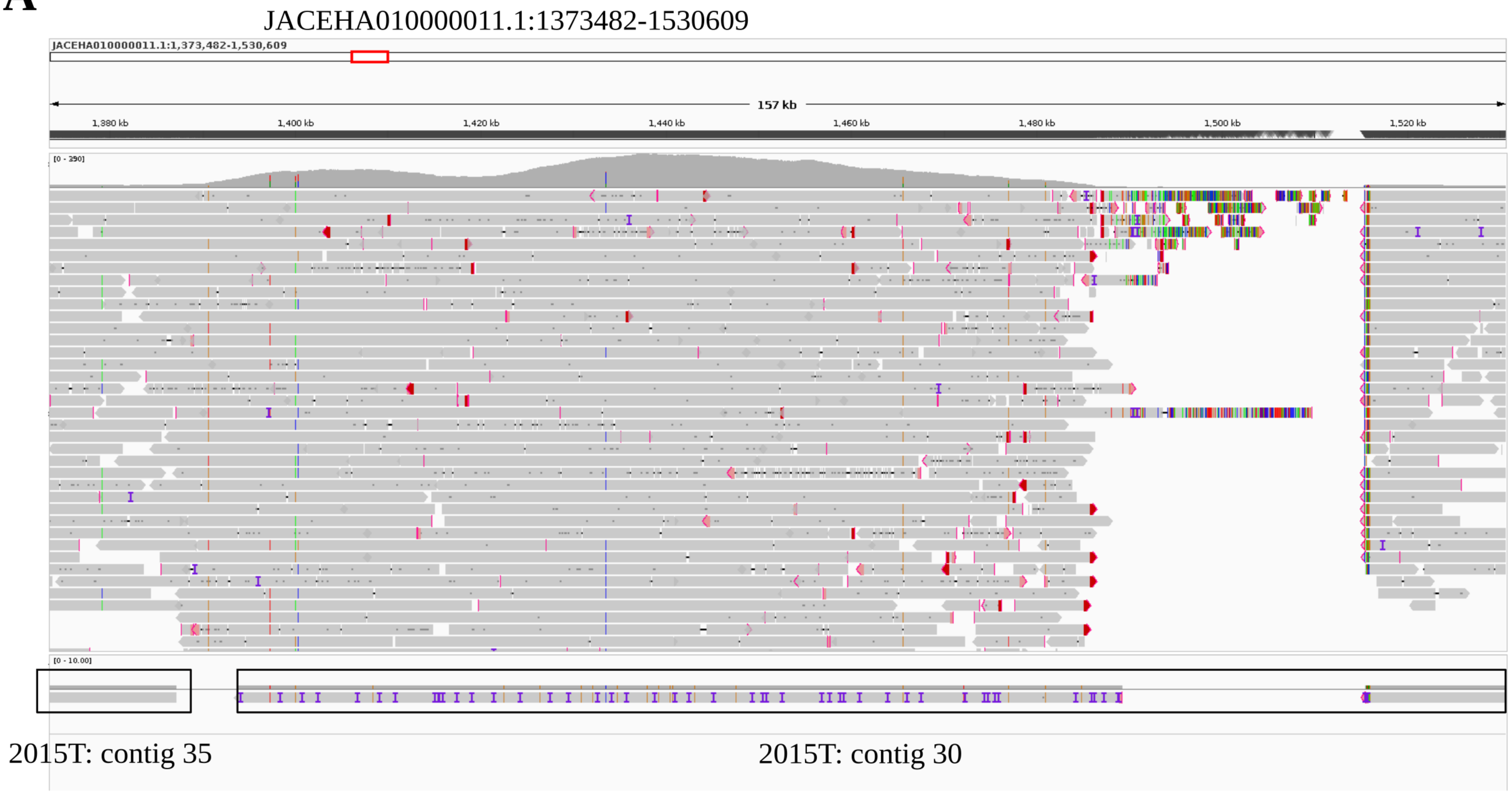

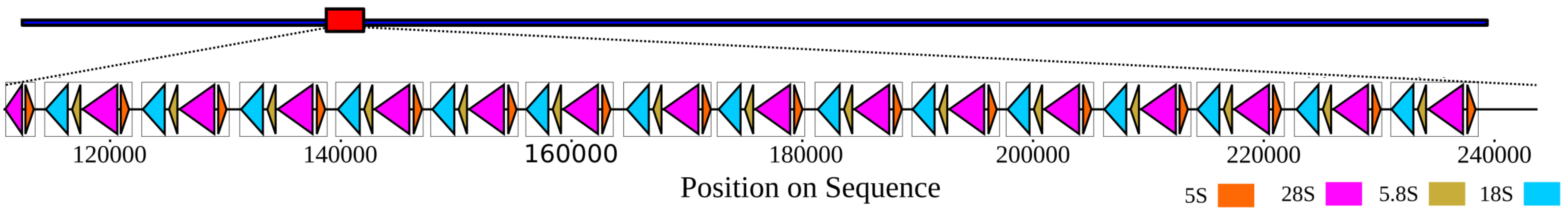

### Supplementary Figure 4

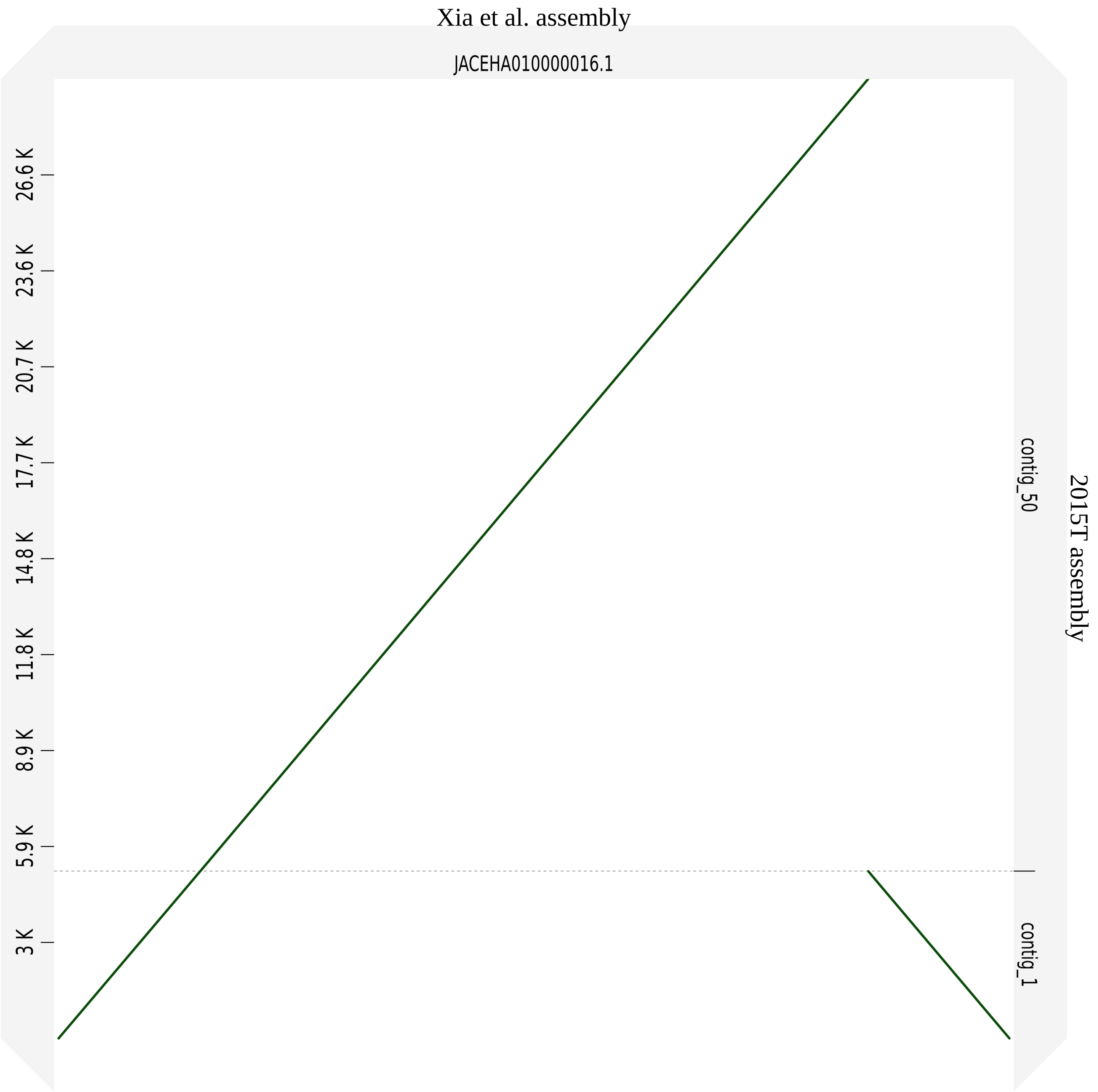

### Supplementary Figure 6

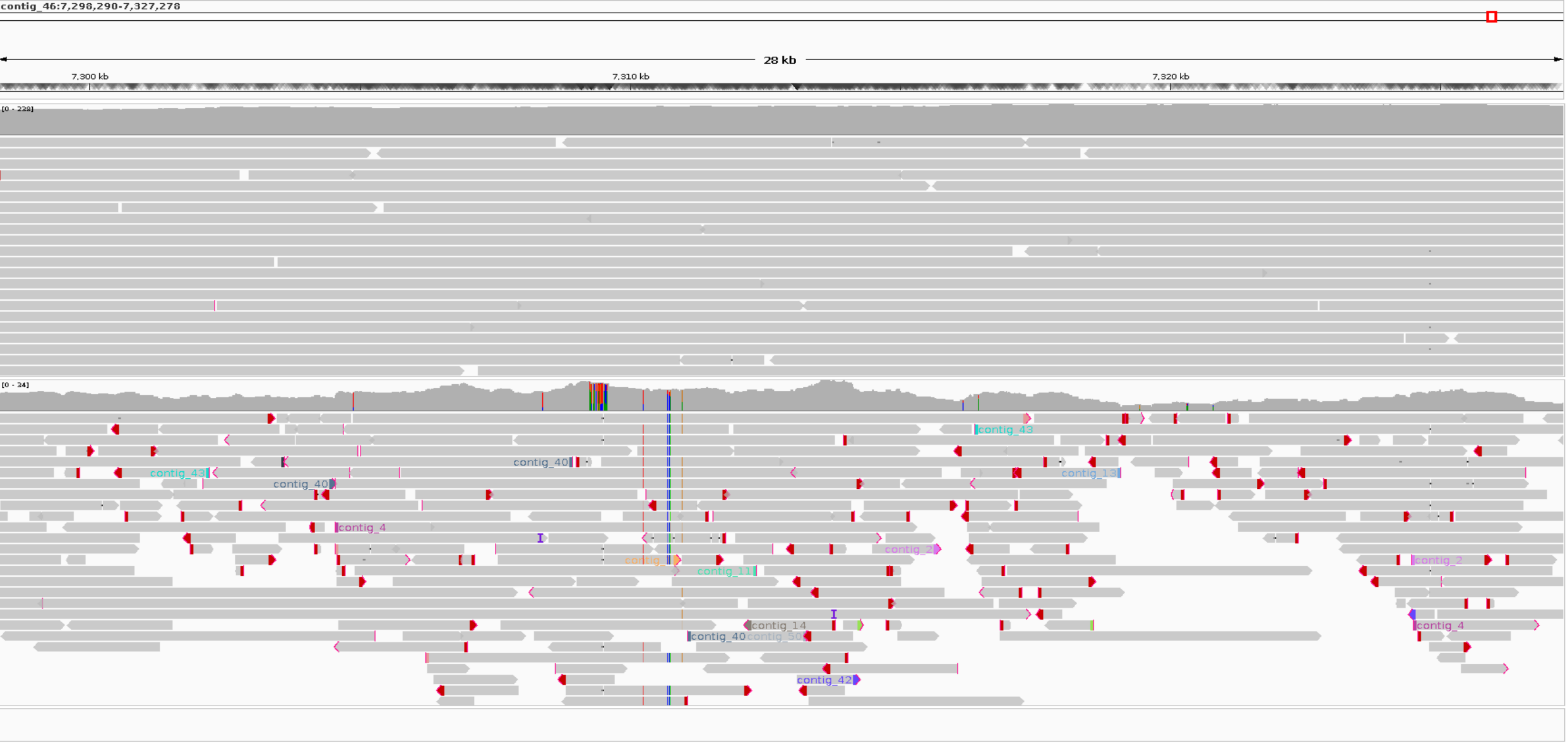

### Supplementary Figure 7

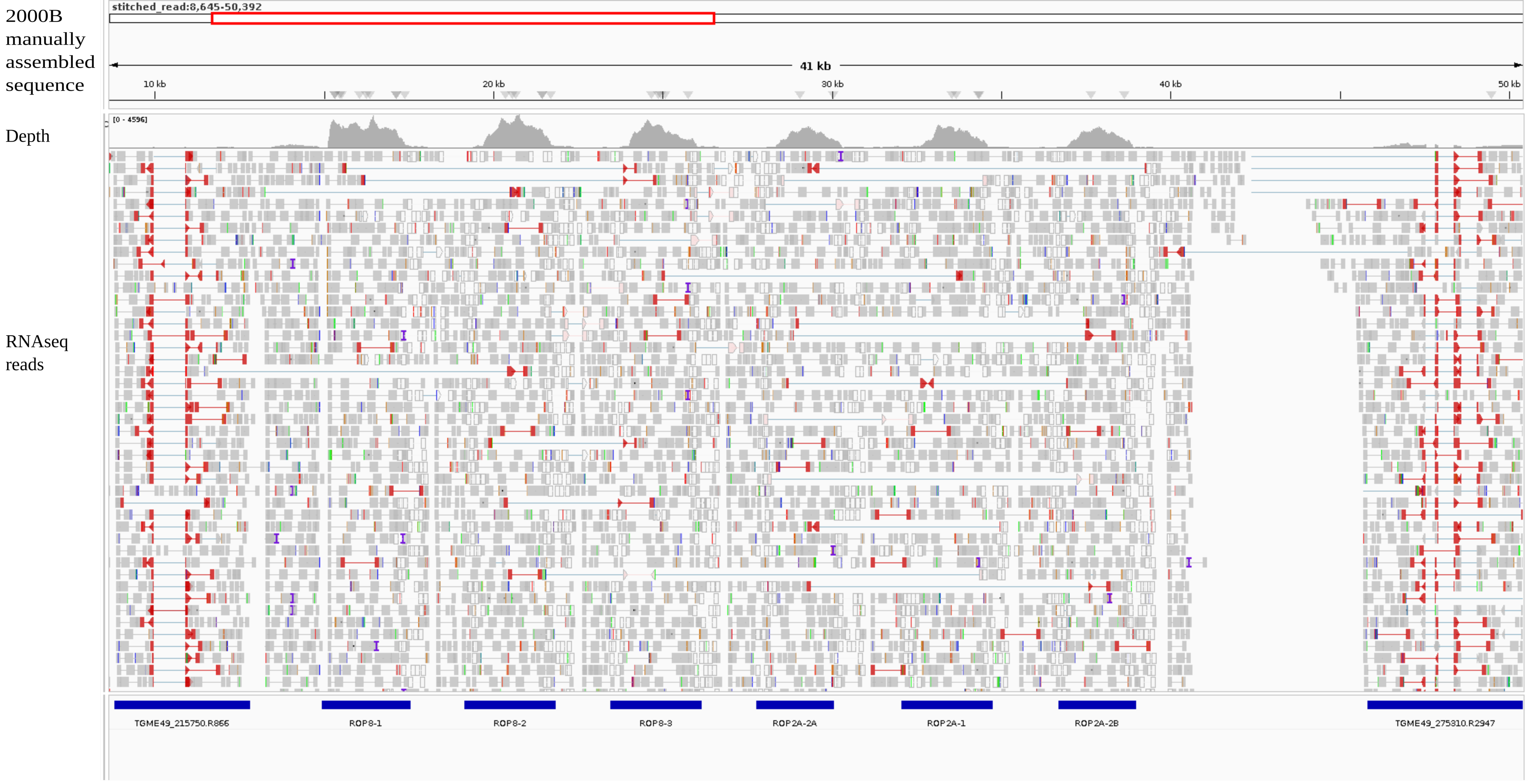

### Supplementary Figure 8

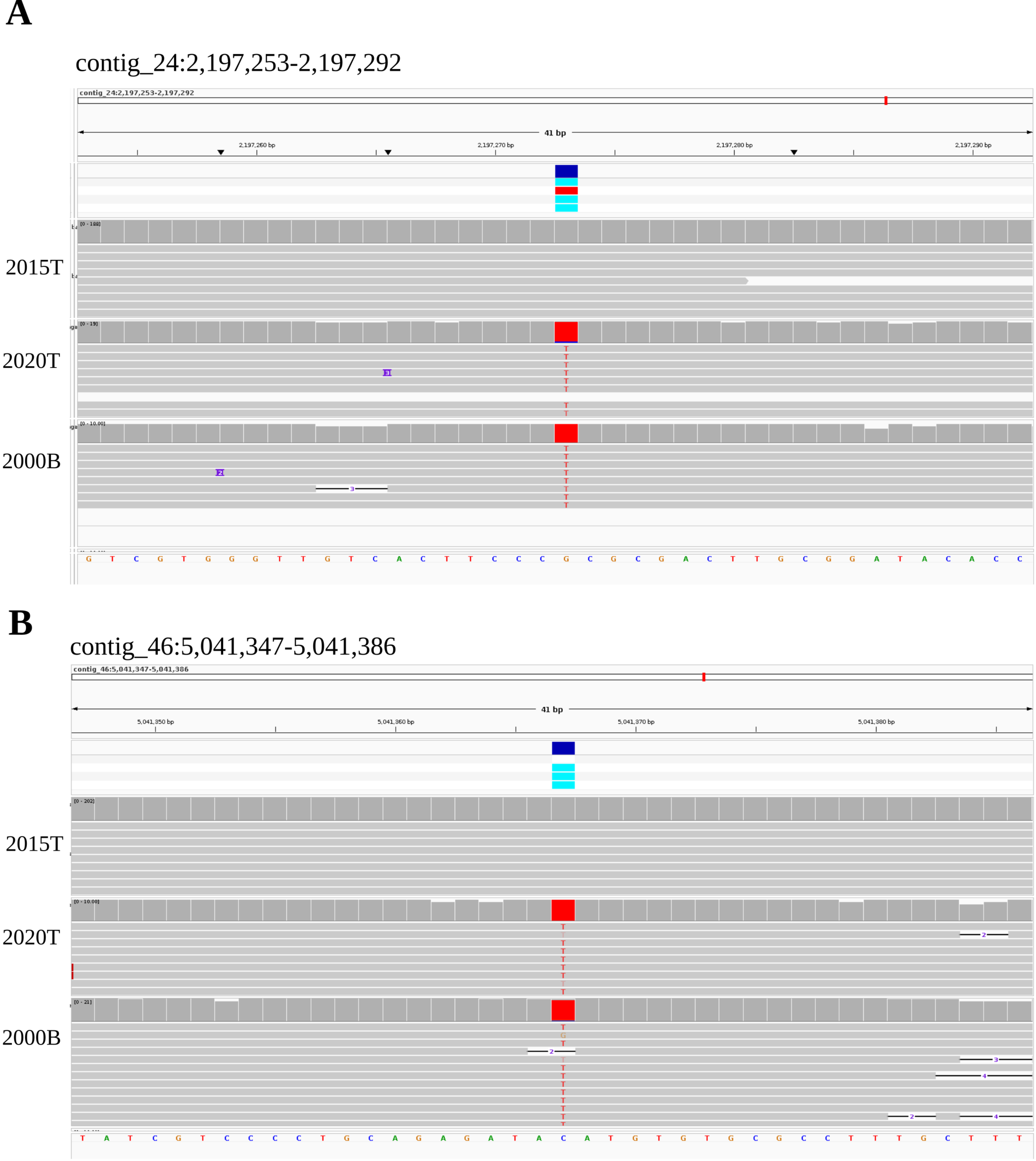

### Supplementary Figure 10

#### Pairwise Exon Similarity for TgME49_IV0036400.2

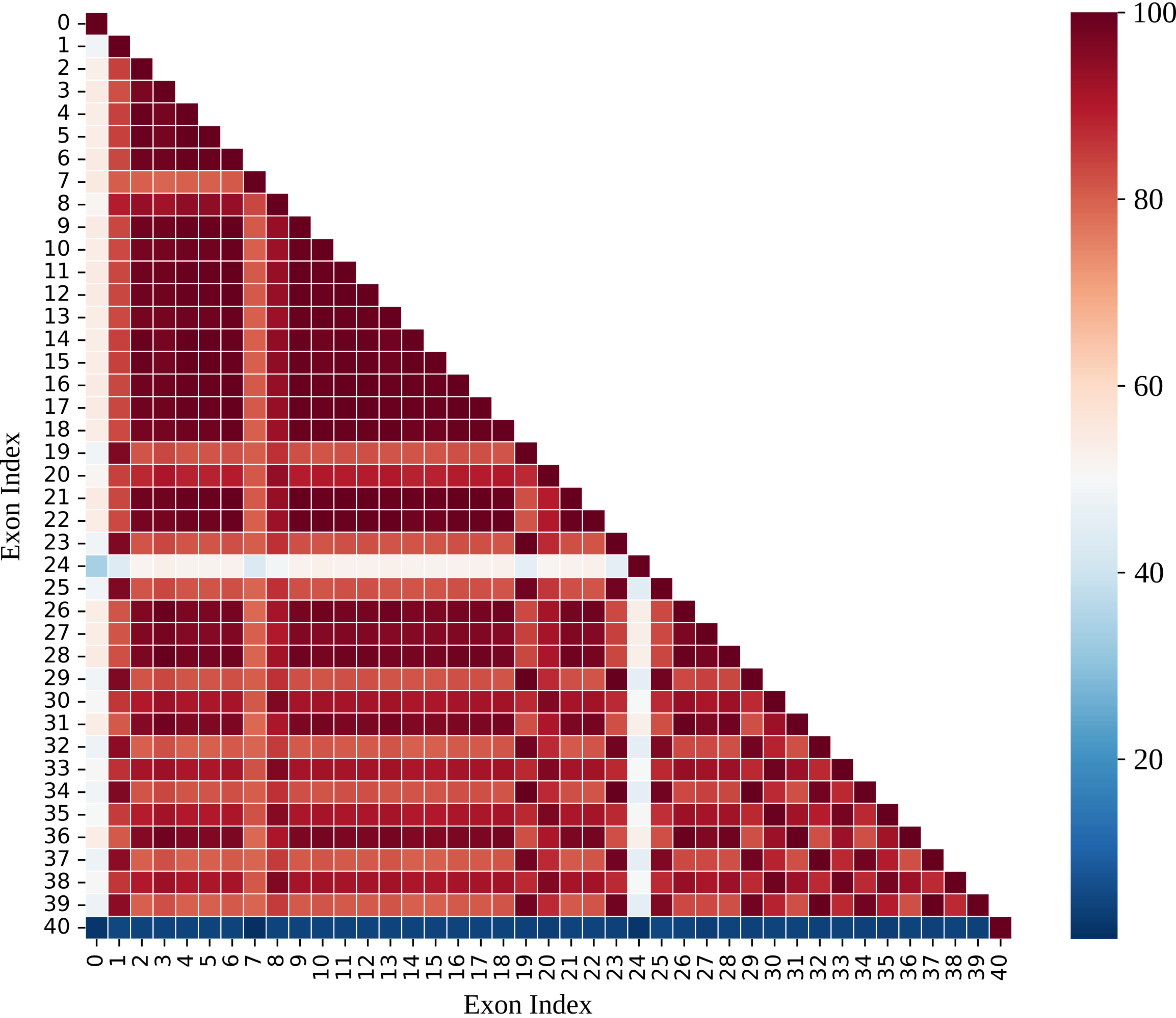
