## Supplementary Figure 2 for "Intra-Strain Genetic Heterogeneity in *Toxoplasma gondii* ME49: Oxford Nanopore Long-Read Sequencing Reveals Copy Number Variation in the ROP8-ROP2A Locus"

**A**

### Xia et al. assembly

### 2015T assembly

THH1 domain-containing protein

JACEHA010000003.1:1,147,141-1,147,630

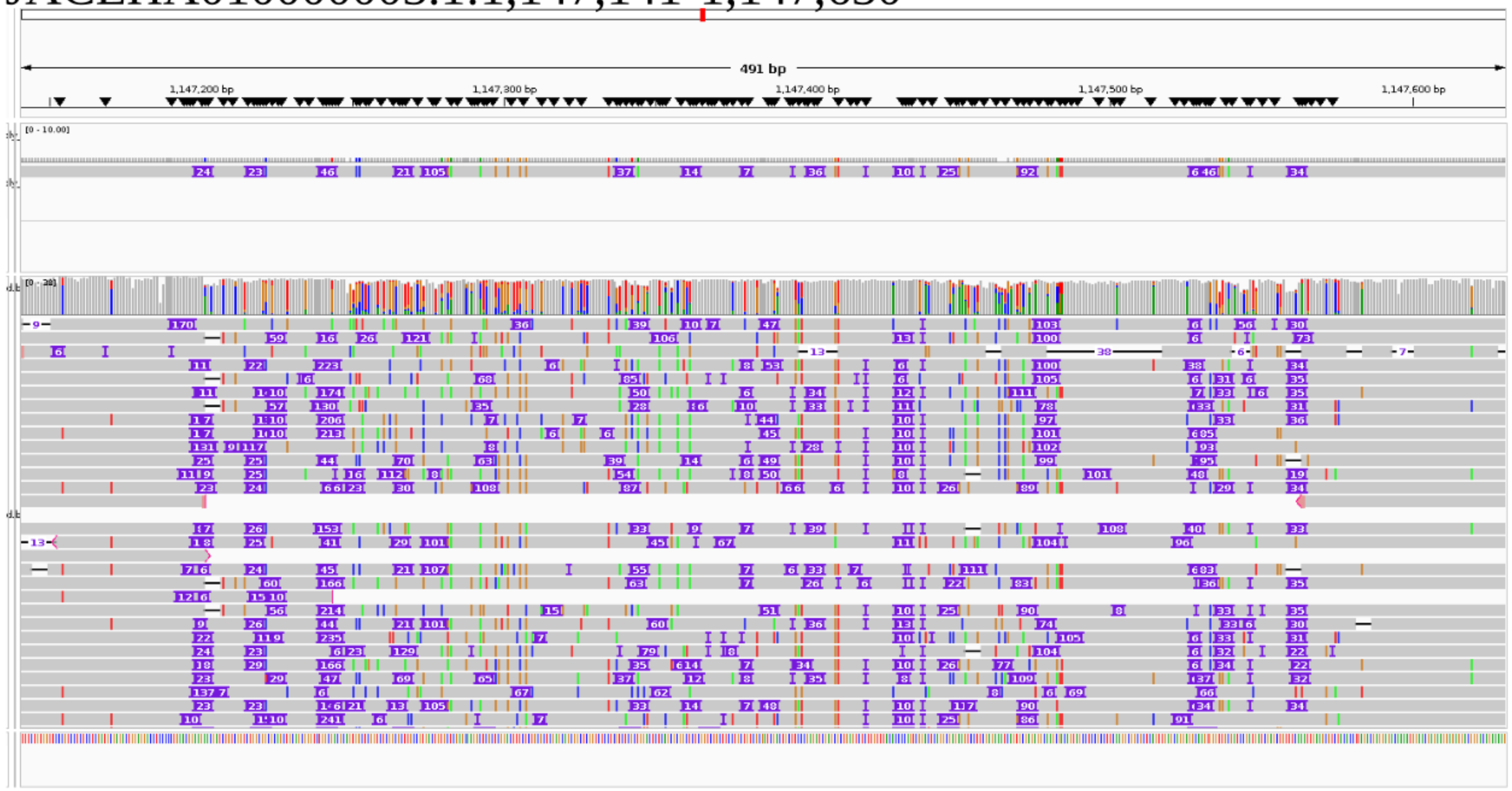


contig_13:1,149,752-1,150,848

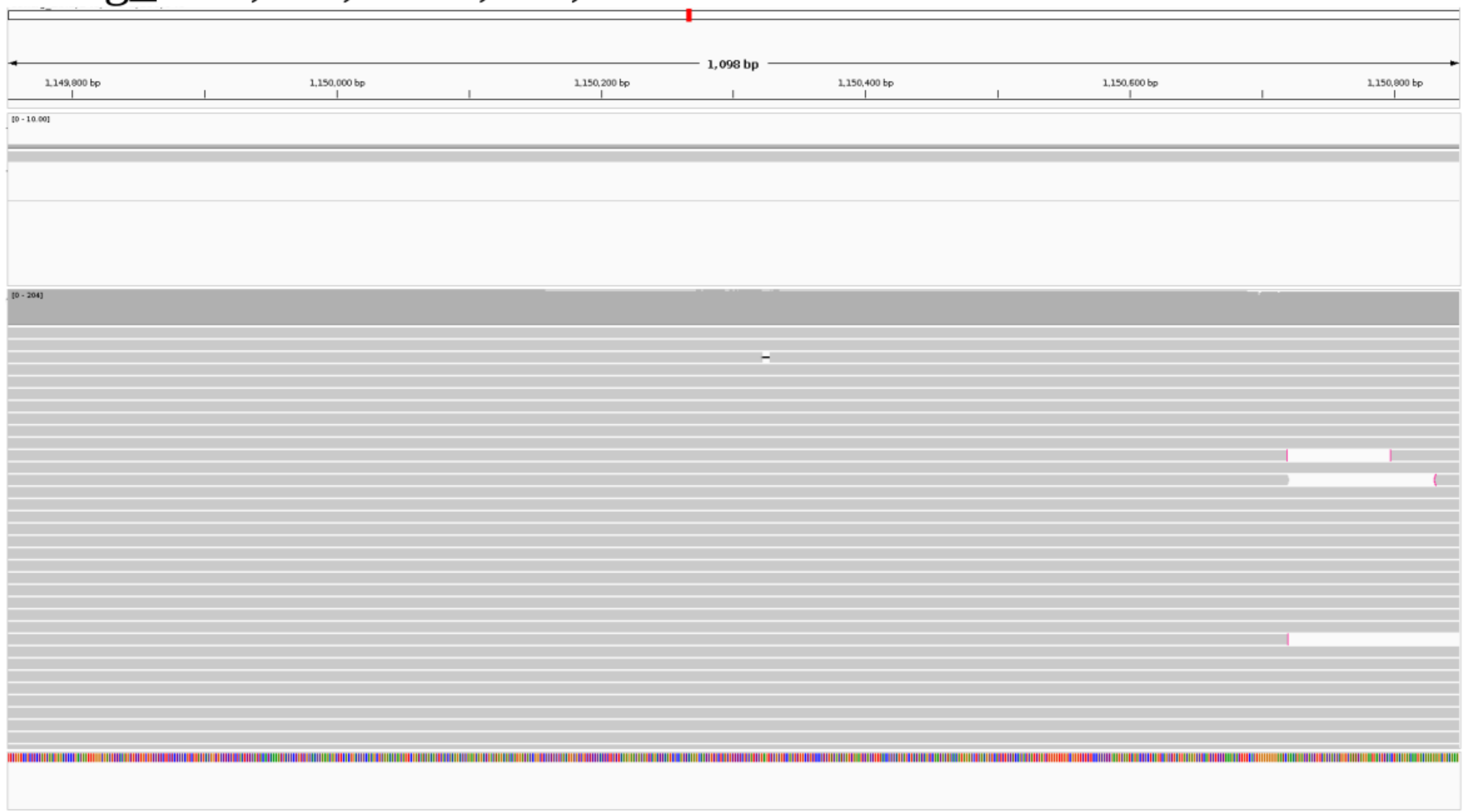


**B**

Malate dehydrogenase MDH

JACEHA010000005.1:1,444,167-1,445,047

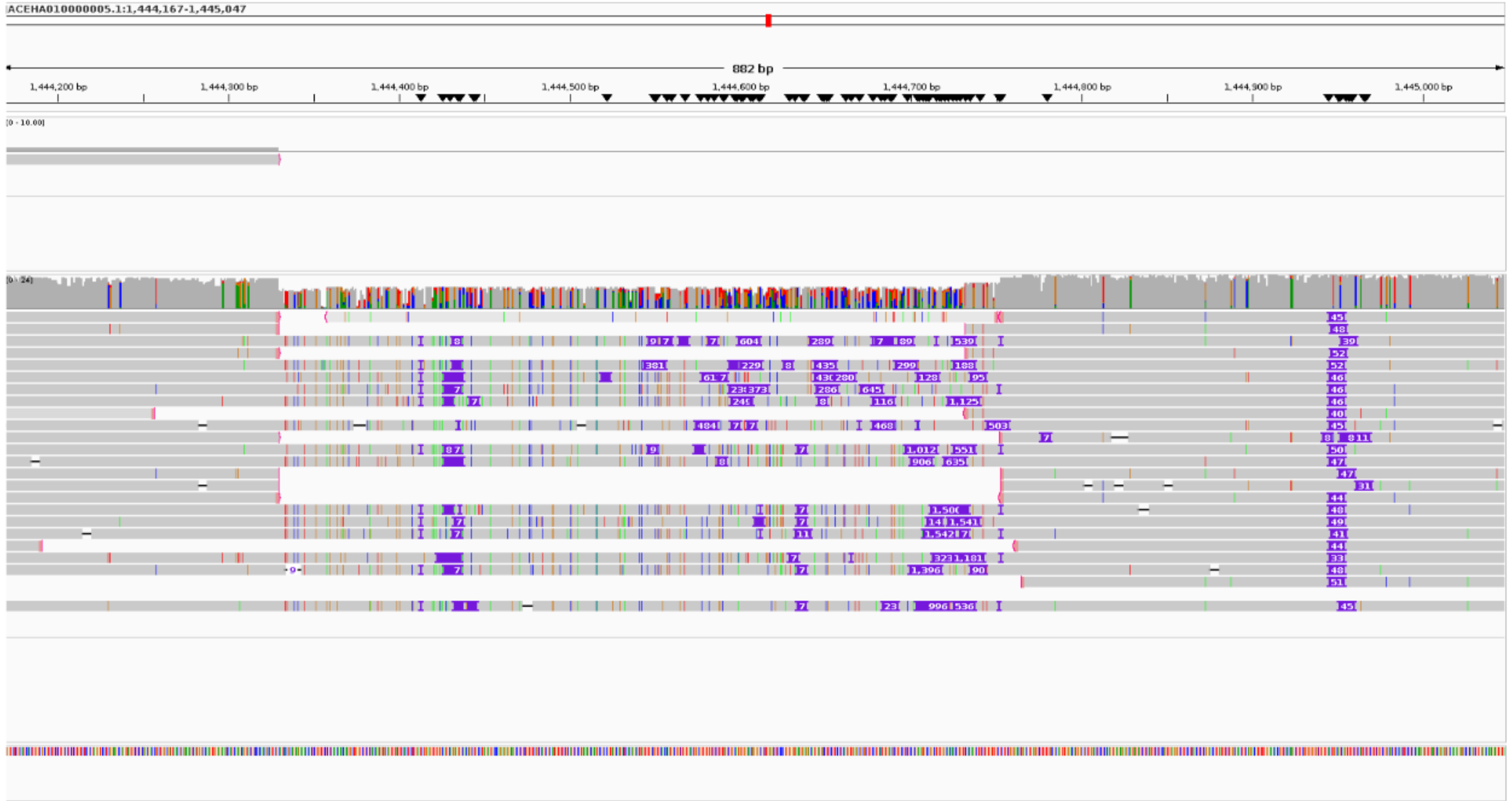


contig_2:1,491,135-1,493,717

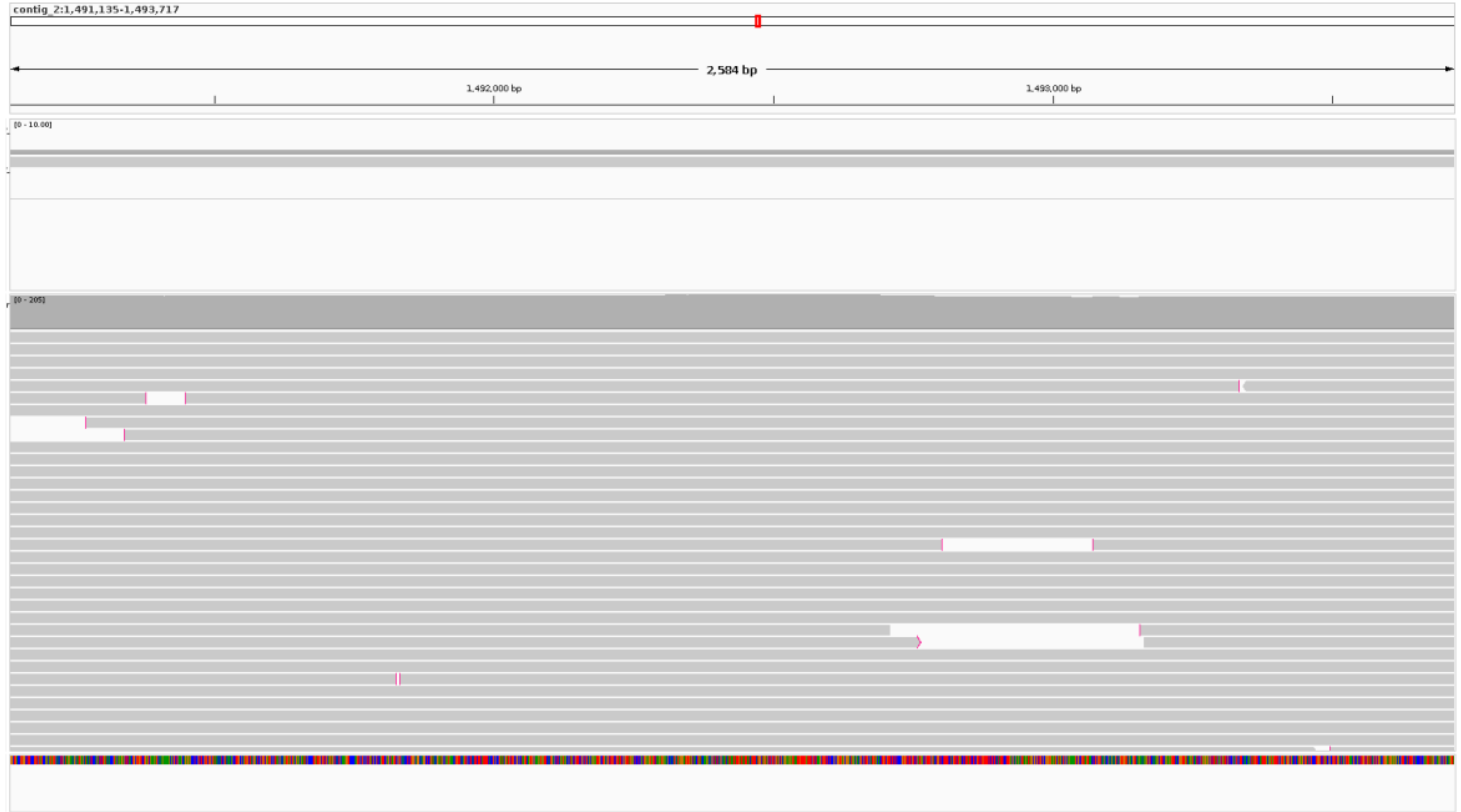
