## Supplementary Figure 5 for "Intra-Strain Genetic Heterogeneity in *Toxoplasma gondii* ME49: Oxford Nanopore Long-Read Sequencing Reveals Copy Number Variation in the ROP8-ROP2A Locus"

**A**

2000B reads against 2015T assembly before modification

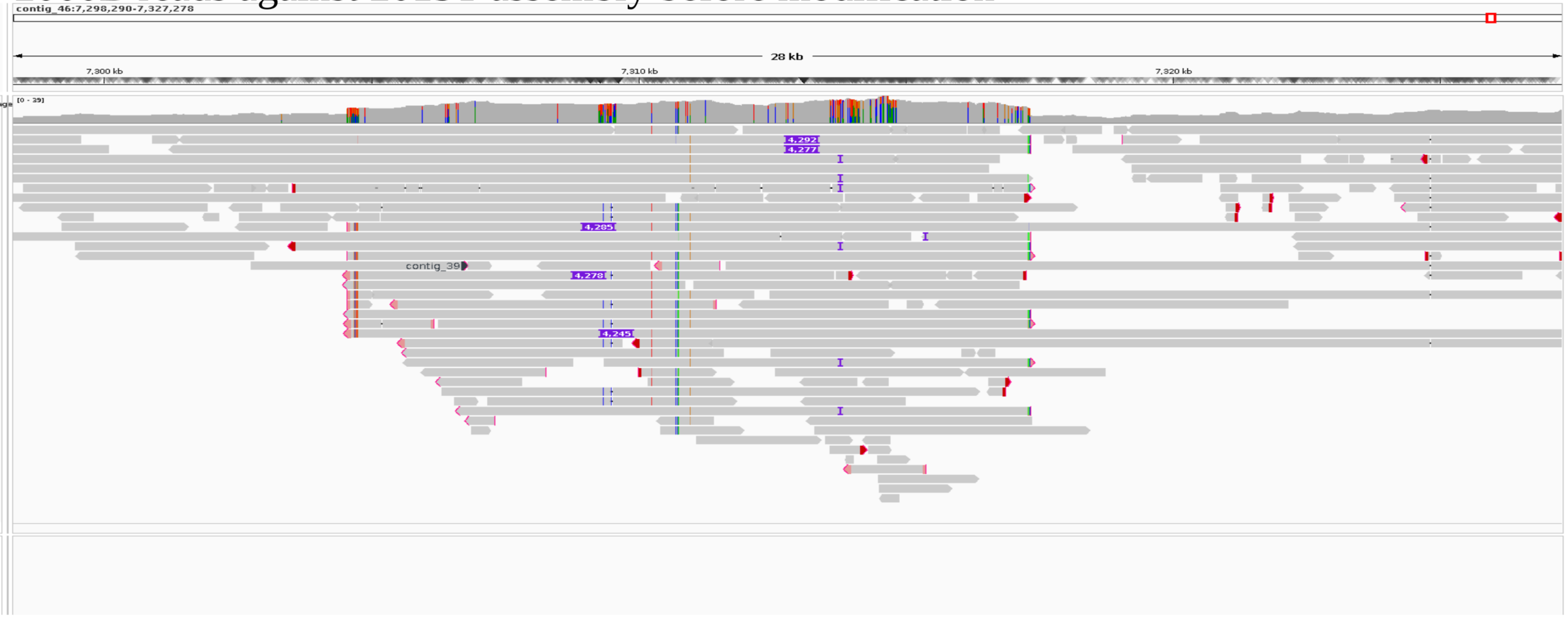


**B**

2000B reads against 2015T assembly after modification

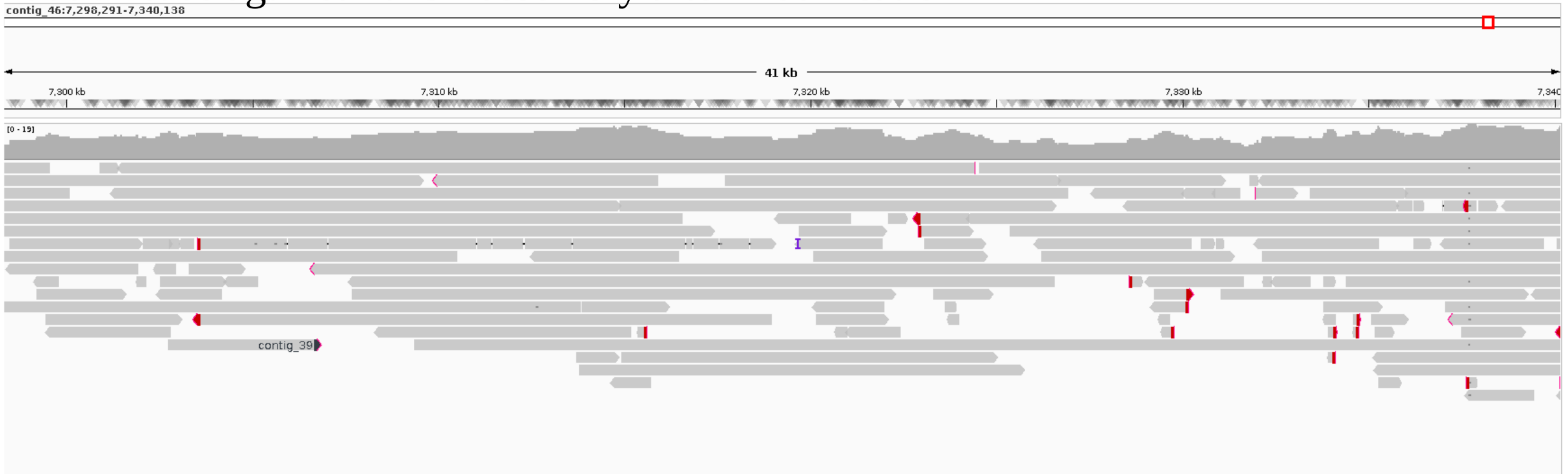
